# RNA Architecture Underlies Discontinuous Transcription and Evolution of Coronavirus

**DOI:** 10.1101/2025.03.13.642953

**Authors:** Zi Wen, Lei Chen, Dehua Luo, Ju Sun, Liangrong Guo, Yingxiang Deng, Zhiyuan Huang, Yuxiang Wang, Ke Pan, Fan Wang, Shaobo Xiao, Li Li, Dengguo Wei

**Affiliations:** National Key Laboratory of Agricultural Microbiology, College of Veterinary Medicine, Huazhong Agricultural University, Wuhan, Hubei, China; Hubei Hongshan Laboratory, Interdisciplinary Sciences Institute, Huazhong Agricultural University, Wuhan, Hubei, China; College of Informatics, Huazhong Agricultural University, Wuhan, Hubei, China; National Reference Laboratory of Veterinary Drug Residues (HZAU) and National Safety Laboratory of Veterinary Drug (HZAU), MOA Key Laboratory for Detection of Veterinary Drug Residues, MOA Laboratory for Risk Assessment of Quality and Safety of Livestock and Poultry Products, Wuhan, Hubei, China; Frontiers Science Center for Animal Breeding and Sustainable Production, Wuhan, Hubei, China; Shenzhen Institute of Nutrition and Health, Huazhong Agricultural University, Shenzhen, China; Shenzhen Branch, Guangdong Laboratory for Lingnan Modern Agriculture, Genome Analysis Laboratory of the Ministry of Agriculture, Agricultural Genomics Institute at Shenzhen, Chinese Academy of Agricultural Sciences, Shenzhen, China; National Key Laboratory of Green Pesticide, International Joint Research Center for Intelligent Biosensor Technology and Health, Central China Normal University, Wuhan, China

## Abstract

Coronaviruses employ discontinuous transcription to produce canonical subgenomic RNAs (sgRNAs) essential for gene expression, as well as non-canonical sgRNAs with unclear function. Nevertheless, the mechanisms regulating the balance between canonical and non-canonical sgRNAs throughout the viral replication cycle remain unknown. We conducted an integrated analysis on 234 transcriptome and 12 RNA-RNA interactome samples across various coronavirus genera. RNA-RNA interactions were identified between TRS-B and TRS-L flanking regions in the same genomic direction, correlating with the formation of canonical junctions. Non-canonical junctions frequently span short or long genomic distance, with short-range junctions generally mediated by reverse complementary stem-loops and coinciding with genomic deletion regions. Conserved long-range non-canonical sgRNAs were identified across different coronaviruses, and these sgRNAs harbor ORF10 or an evolving gene to suppress antiviral innate immune responses. This work establishes a structural framework that enables an understanding of the regulatory mechanism behind discontinuous transcription and the evolutionary pathway of coronaviruses.

**Teaser:** Deciphering coronavirus transcription and evolution: an integration analysis of transcriptome and RNA-RNA interactome.

**Graphical Abstract:** 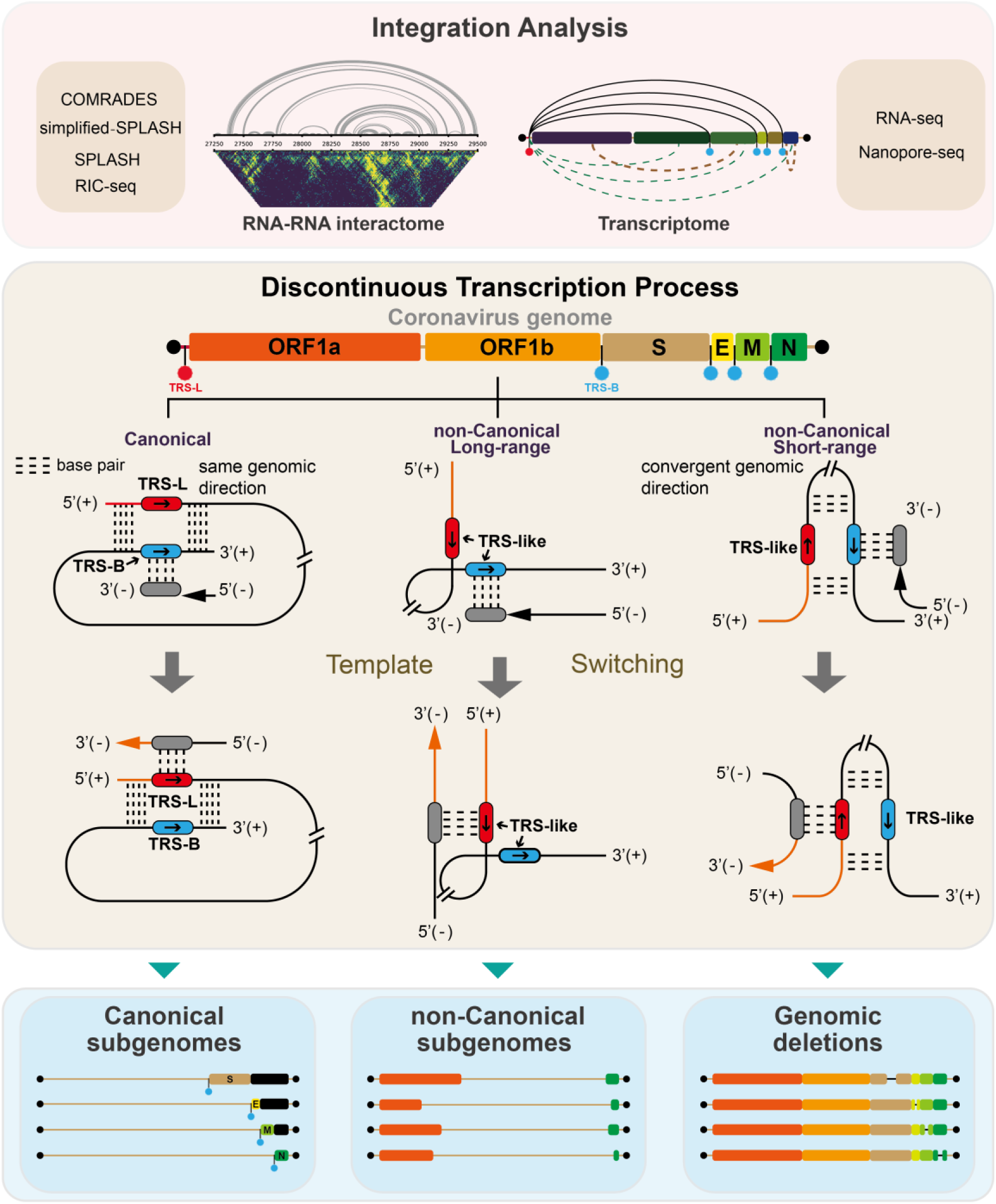

---

Coronaviruses, such as SARS-CoV-2, utilize a discontinuous transcription strategy to produce subgenomic RNAs (sgRNAs), a process crucial for the regulation of gene expression and evolutionary adaptation (*1–5*). Canonical sgRNAs are generated through a complex template-switching mechanism in which the RNA-dependent RNA polymerase (RdRp) dynamically switches templates from the transcription regulatory sequence in the body (TRS-B) to that in the leader (TRS-L) during RNA synthesis (*6*). The efficiency of the discontinuous transcription depends not only on the conservation between TRS-L and TRS-B, but also on the rearrangement of the genome into a conformation suitable for template-switching (*6–8*). However, the precise mechanisms and details of this process remain to be fully elucidated. Besides canonical sgRNAs, recent studies have identified TRS-independent non-canonical sgRNAs in SARS-CoV-2 (*2*, *3*, *9*, *10*), further highlighting the complexity of discontinuous transcription. We noticed that the proportion of canonical and non-canonical junctions varied at different phases of viral replication, but the regulatory mechanism is unknown. Deciphering RNA-RNA interactions within the viral genome is essential for identifying drivers of template switching and for understanding the roles of multiple sgRNAs.

Advances in transcriptome and RNA-RNA interactome sequencing techniques have offered new tools to study the discontinuous transcription mechanism. RNA-seq data provides precise short sequencing reads, facilitating the identification of junction sites within sgRNAs (*7*). Nanopore sequencing allows the comprehensive identification of sgRNAs and enables accurate quantification of their transcription levels (*3*). Genome-wide RNA-RNA interactions within the coronavirus have been inferred by methodologies such as COMRADES (*11*), SPLASH (*12*), simplified-SPLASH (*13*), and RIC-seq (*14*), which rely on capturing proximity-linked chimeric reads. The abundance of these chimeric reads indicates the RNA-RNA interaction strength (*14*), with stronger interactions suggesting closer spatial proximity. This significant interaction strength can thus serve as predictive constraints for base pairing within RNA architectures, which are intricately linked to transcription and replication (*11–14*). Zhang et al. identified distant interactions between TRS-L and TRS-B in the SARS-CoV-2 genome through simplified-SPLASH, suggesting a potential role of genome architecture in mediating canonical sgRNA formation (*13*). However, the specific RNA architecture affecting the formation of canonical sgRNAs in coronaviruses has not been systematically characterized, and the architecture as well as the potential roles of non-canonical sgRNAs remain unclear, warranting further comprehensive investigation.

Coronaviridae family comprises four genera: α-, β-, γ-, and δ-coronaviruses. Current genome-wide RNA-RNA interaction analyses primarily focus on β-coronaviruses, such as SARS-CoV-2 and MERS-CoV. In this study, we performed RNA-seq and RIC-seq experiments on the α-coronavirus PEDV (Porcine Epidemic Diarrhea Virus) and the δ-coronavirus PDCoV (Porcine deltacoronavirus). Utilizing public and our own data, we compiled a dataset of 234 transcriptomes and 12 RNA-RNA interactomes from four coronavirus genera and processed it for storage in the Coronavirus Omics Data Explorer (CODE: https://webofgroup.cn/dgwei/hngene/#/index). Our findings show that the ratio of canonical to non-canonical junctions is influenced by extrinsic factors, with each type exhibiting distinct RNA architectures. Short-range non-canonical junction sites coincide with stem-loops, potentially facilitating the generation of genome-defective viral strains. Notably, conserved long-range non-canonical sgRNAs are identified to harbor a gene or an evolving gene in suppressing the antiviral innate immune response. This study provides valuable insights into the mechanisms governing viral transcription and evolution at the RNA architecture level.

## Results

### Integration of transcriptome and RNA architecture profiles

To explore the influence of genome architecture to transcription, we have compiled a dataset comprising 12 RNA architecture profiles and 234 transcriptome data, including 165 RNA-seq samples and 69 nanopore samples, from 4 genera of coronaviruses (Fig. 1A). The dataset procured in this work was processed and archived in CODE, a repository we developed and made accessible online (Fig. 1B). RNA-seq and nanopore sequencing data were employed to detect the types and quantities of sgRNAs, providing the transcriptomic landscape from different host cells upon coronavirus infection. Additionally, the complete sgRNA information derived from nanopore sequencing reads enables measurement of the transcription level of each viral gene (Fig. 1A).

**Figure 1.**
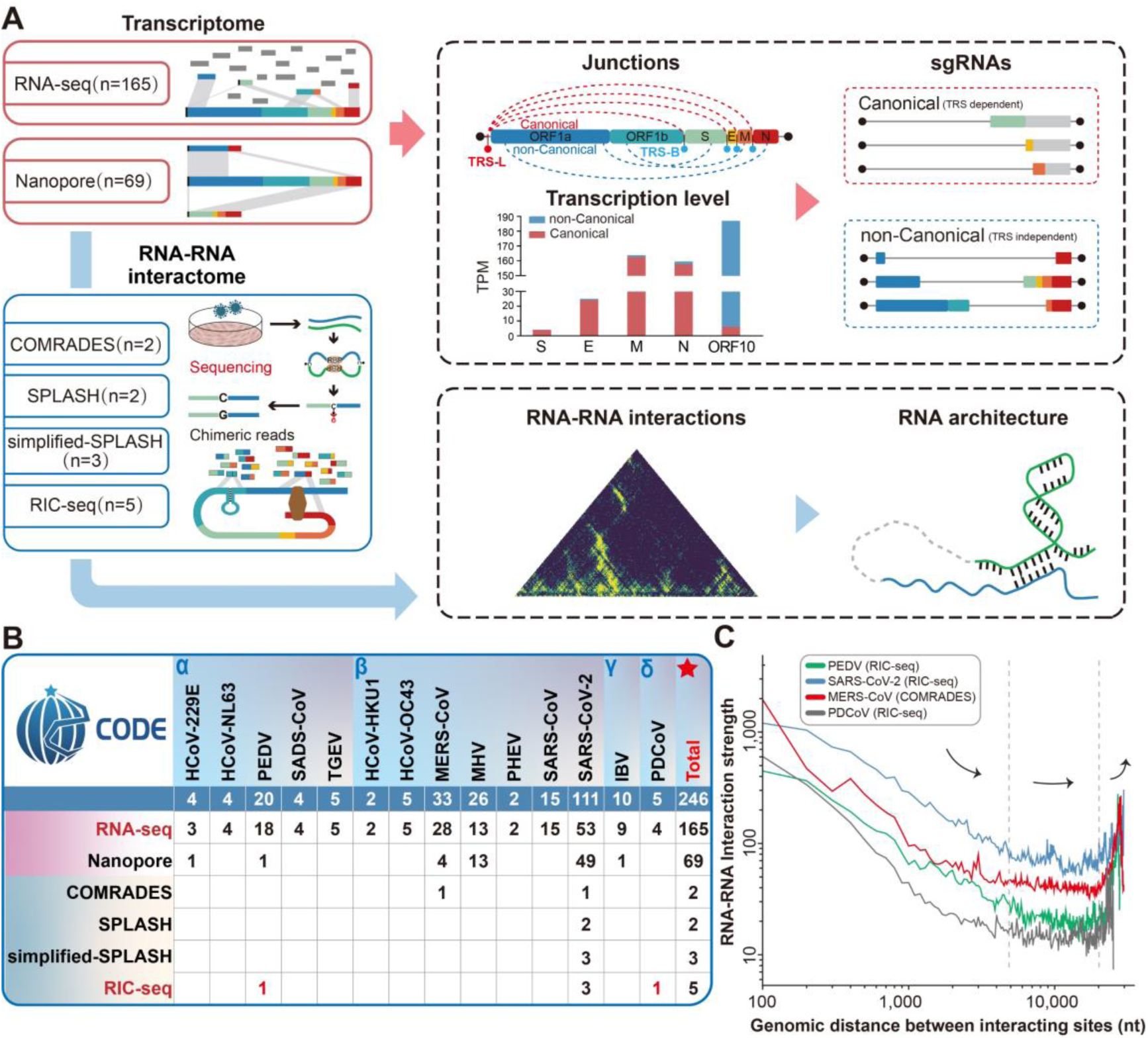
Integration analysis for RNA-RNA interactome and transcriptome profiles. (A) Schematic of the integrated analysis of RNA-RNA interactome and transcriptome data. Through the identification of RNA structures and junction sites within sgRNAs, we endeavor to comprehensively analyze the association between genome architecture and transcription in coronaviruses. (B) A comprehensive compilation of transcriptome and RNA-RNA interactome data for 14 coronaviruses. The RIC-seq data of PEDV and PDCoV are generated in this work, with corresponding RNA-seq data supplemented for both PEDV and PDCoV. (C) RNA-RNA interaction strength across a range of distances between pairwise interacting RNAs from different coronaviruses.

Based on genome-wide RNA-RNA interactions captured through proximity-linked chimeric reads, we can infer the global or local architecture of the coronavirus genome (Fig. 1A). Public architecture profiles, including data obtained through COMRADES, SPLASH, simplified-SPLASH, and RIC-seq, are primarily focused on β-coronaviruses SARS-CoV-2 and MERS-CoV (Fig. 1B). We probed RNA-RNA interactions from cells infected with PEDV and the PDCoV utilizing RIC-seq and supplemented these with RNA-seq data for each virus (Fig. S1). A consistent ‘U-shaped’ distribution of genome-wide RNA-RNA interactions is observed across various coronaviruses and experimental methodologies (Fig. 1C and Fig. S2A). Within a genomic span of approximately 5,000 nt, a rapid decline in the interactions is observed as the distance increased, exhibiting a logarithmic slope close to -1 (Fig. 1C). Interactions reached a minimum between genomic distances ranging from 5,000-20,000 nt, and a gradual increase in interactions is observed beyond approximately 20,000 nt (Fig. 1C). These results indicate a prevalence of short- and long-range genomic interactions, with a relative lack of mid-range genomic interactions.

### The proportion of non-canonical junctions might act as a biomarker indicative of viral transcription activity in infected cells

We further selected 103 RNA samples that were manually validated to have a virus genome mapping rate greater than 0.5% and more than 5,000 discontinuous transcription events. These samples, which encompass 14 different coronaviruses, were obtained under various conditions including different passages, hours post-infection (hpi), infected cell types, multiplicity of infection (MOI), and with/without antiviral drugs. The analysis of discontinuous transcriptional events reveals that canonical junctions are prevalent in the majority of the RNA-seq datasets (Fig. 2A). For instance, more than 50% of the junctions are canonical in 23 out of 31 SARS-CoV-2 samples and 16 out of 20 MERS-CoV samples (Fig. S2BC). The average proportion of canonical junctions across all RNA-seq samples is 69.07% (Fig. S2D).

**Figure 2.**
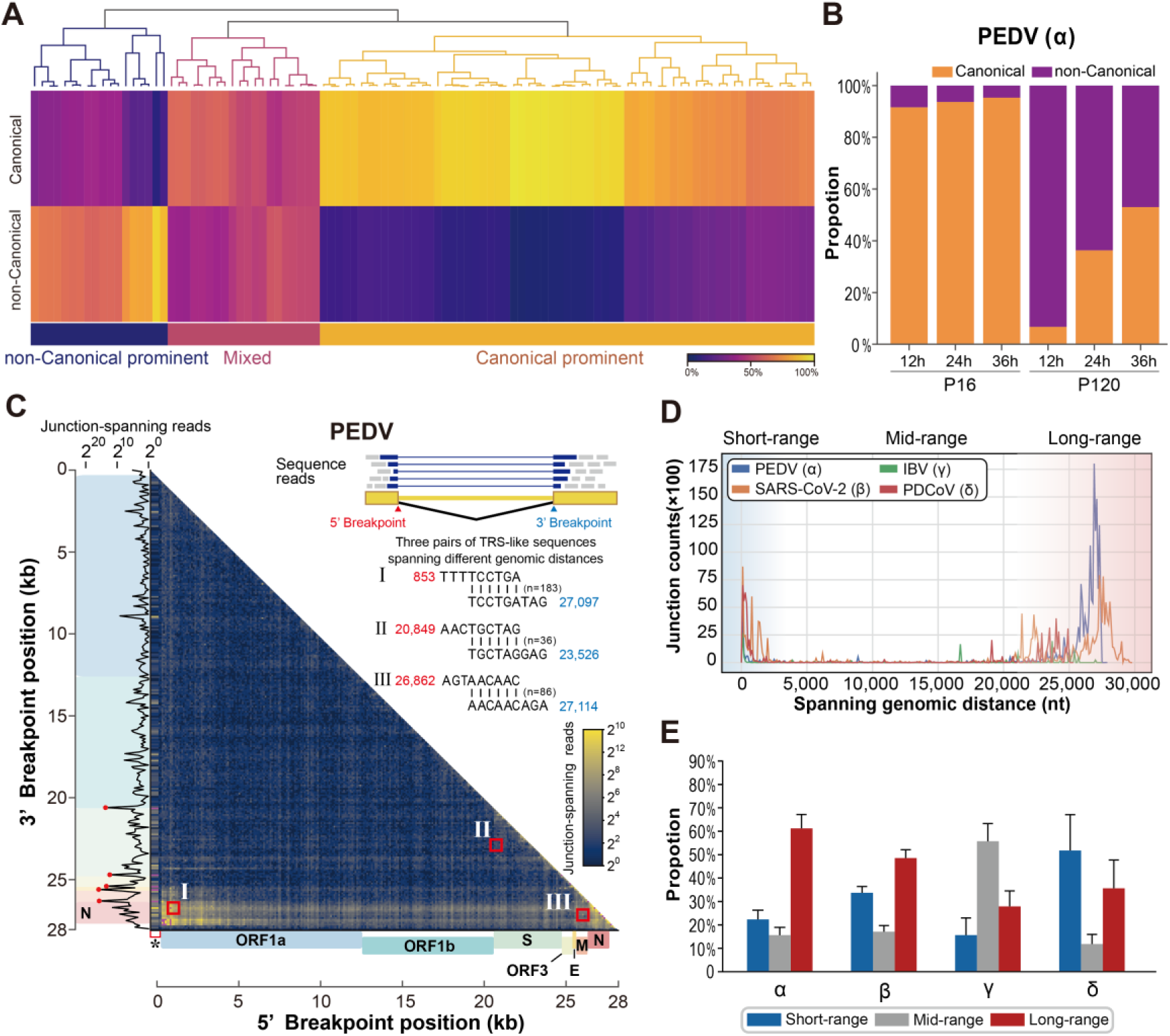
Characterization of non-canonical junctions. (A) Hierarchical clustering for the proportion of non-canonical and canonical junctions based on 103 RNA-seq samples. (B) Proportion of canonical and non-canonical junctions in RNA-seq samples from porcine small intestinal 2-D enteroid cells infected with PEDV. These samples were infected with low-passage (16 passages, P16) and high-passage (120 passages, P120) PEDV strain and detected at 12, 24 and 36 hpi. (C) The heatmap depicts the distribution of junction sites for PEDV genome. Gene annotations are displayed on the bottom and left sides of the heatmap. The line plot on the left side illustrates the distribution of 3’ breakpoints of chimeric reads. The top right shows the three pairs of TRS-Like sequences involved in non-canonical junctions at different genomic distances. (D) Visualization of the distribution of non-canonical junctions at different spanning genomic distances from 4 genera of coronaviruses. Non-canonical junctions predominantly occur in short-range and long-range genomic distance. (E) Average proportion of non-canonical junctions with different genomic distance across 4 genera of coronaviruses.

Non-canonical junctions predominant in approximately one quarter of all RNA-seq samples. For example, 3 out of the 4 MERS-CoV samples with predominantly non-canonical junctions were collected from Vero E6 cells infected at 72 hpi (Fig. S2C). Interestingly, among these RNA-seq samples obtained from different passages of PEDV (*15*), 93.58% of junction events are canonical in the low-passage (16 passages, P16) samples, whereas non-canonical junction events predominate in the high-passage (120 passages, P120) samples (Fig. 2B). In addition, the proportion of non-canonical junction decreases as hpi increases in both P16 and P120 samples (Fig. 2B). These findings exhibit that the proportions between canonical and non-canonical junction events are dynamic across different samples (Fig. 2A), suggesting that non-canonical junctions may play a critical role in viral transcription under certain conditions.

### Characterization of non-canonical discontinuous transcription events

We constructed the transcriptional landscape of cells infected with PEDV and PDCoV through RNA-seq (Fig. S1A). Both canonical and non-canonical junctions were identified in comparable numbers from Vero cells infected with PEDV, including 144,832 canonical junctions between pairs of TRS and 159,290 non-canonical junctions involving TRS-independent sequences (Fig. 2C). Genomic RNA pairs involved in the formation of non-canonical junctions exhibit higher consistency across different coronavirus genomes compared to control (Fig. S3). Figure 2C depicts three pairs of identical sequences involved in non-canonical junctions at varying genomic distances. These sequence pairs with high consistency, independent of TRS, are here defined as TRS-like sequences.

Across different coronavirus genomes, non-canonical junctions are predominantly identified between the N gene and ORF1a over extended genomic distances, as well as between pairs of structural protein encoding regions at close genomic distances (Fig. 2C and Fig. S4A). Specifically, our PEDV RNA-seq sample reveals 146,363 long-range junctions (> 22,000 nt), comprising 91.88% of total non-canonical junctions, in addition to 7,944 short-range junctions (< 1,700 nt, 4.99%) and 4,992 mid-range junctions (1,700 - 22,000 nt, 3.13%) (Fig. 2D). By classifying based on genomic distances, non-canonical junctions in other coronaviruses could also be categorized into three distinct groups (Fig. 2D, Fig. S4A and Table S1). The average proportion of short-range, mid-range and long-range junctions is 31.46%, 18.70% and 49.84% among all RNA-seq samples (Fig. S4B). Consistently, long-range and short-range junctions are the predominant types of non-canonical junctions among α-, β-, and δ-coronaviruses, with mid-range junctions constituting a minor fraction of these events (Fig. 2E).

### TRS-B and TRS-L flanking regions exhibit RNA-RNA interaction with same genomic direction

Combination analyses of transcriptome and RNA-RNA interactions data were performed for 4 coronaviruses, with these two types of data for each virus obtained under the same experimental conditions of infected cells and hpi. Utilizing RIC-seq data for PEDV, SARS-CoV-2, and PDCoV, as well as COMRADES data for MERS-CoV, we generated RNA-RNA interaction heatmaps for the TRS-L and TRS-B flanking regions (encompassing ±50 nucleotides around junction sites). These heatmaps exhibit a notable enrichment along the diagonal for each gene across these coronaviruses (Fig. 3A). With further aggregate peak analysis (APA) (*16*) on interaction profiles of all canonical junction sites, parallel diagonal alignments are noted across diverse coronavirus genomes (Fig. 3B, Fig. S5AB and Fig. S6-S9), which indicate these interactions with same genomic direction. Moreover, a similar phenomenon is observed in the APA heatmaps of canonical junctions for SARS-CoV-2, as detected by other methods such as COMRADES, SPLASH, and simplified-SPLASH (Fig. S5CD).

**Figure 3.**
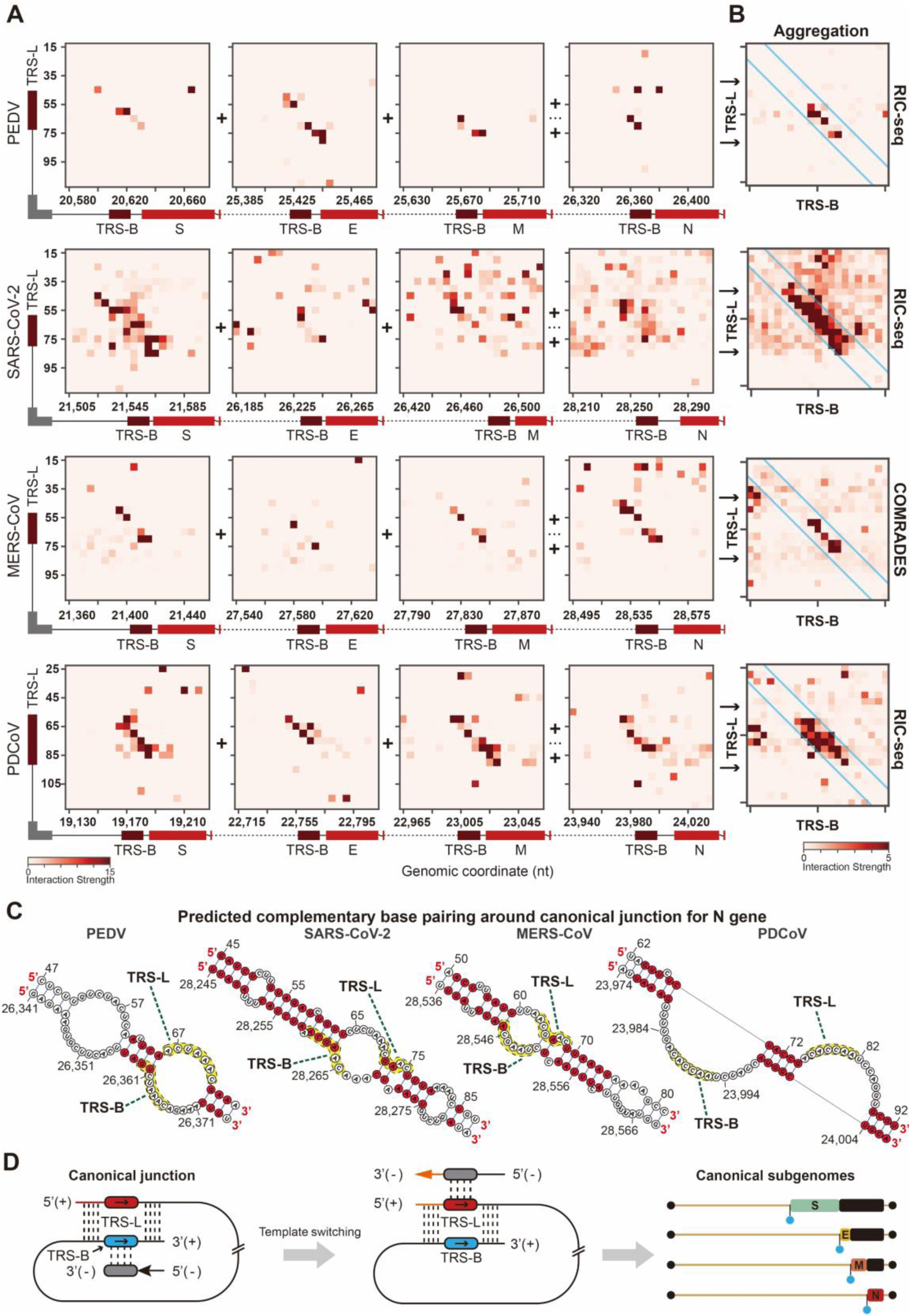
Conserved RNA-RNA interactions patterns under canonical junction. (A) Heatmaps illustrating genomic interactions between the TRS-B and TRS-L flanking regions across various coronavirus genomes. Genome-wide interactions captured by RIC-seq for PEDV, SARS-CoV-2 and PDCoV and by COMRADES for MERS-CoV were used as input. (B) Aggregation peak analysis (APA) plot shows the aggregate RNA-RNA interactions from canonical junction site-flanking regions of PEDV, SARS-COV-2, MERS-CoV and PDCoV. RNA-RNA interactions for PEDV, SARS-CoV-2 and PDCoV were detected by RIC-seq, and RNA-RNA interactions for MERS-CoV were detected by COMRADES. The emphasized parallel diagonal patterns in the APA heatmap signify RNA-RNA interactions that are in same genomic direction. (C) Prediction of complementary base pairings between TRS-L flanking region and TRS-B flanking region of N gene are conducted for each coronavirus, utilizing RNA-RNA interactions with consistent direction as a constraint. (D) The model for canonical sgRNA formation: complementary base pairings between the TRS-L and TRS-B flanking regions bring a pair of TRS into spatial proximity; this alignment facilitates RdRp template switching from TRS-B to TRS-L during sgRNA template synthesis; ultimately, canonical sgRNAs are synthesized by transcription from these templated strands.

Subsequently, RNA-RNA interactions with consistent direction were employed as a constraint to predict complementary base pairings between TRS-L and TRS-B flanking regions. Taking TRS-B flanking region of N gene as an example, these parallel complementary base pairings supported by RNA-RNA interaction are highlighted in red (Fig. 3C), as well as base pairings for other genes (Fig. S6-S9). Notably, the quantity of canonical junctions in genomes of PEDV, SARS-CoV-2 and PDCoV are positively correlated with the number of RNA-RNA interaction-supported base pairings between TRS-L and TRS-B flanking region (Fig. S9B), emphasizing the role of RNA architecture in facilitating canonical junction formation. We propose that these RNA-RNA interactions with same genomic direction contribute to spatial proximity between TRS-L and TRS-B, which promote template switching of RdRp during sgRNA synthesis (Fig. 3D).

### Short-range non-canonical junction sites show overlap with reverse complementary RNA stem-loops

On the genome-wide scale, short- and long-range non-canonical junctions are more prevalent than mid-range junctions (Fig. 2D), a pattern mirrored in the genome-wide RNA-RNA interactions (Fig. 1C), suggesting a potential association between the non-canonical discontinuous transcription events and RNA architectures. We then investigated the RNA-RNA interaction patterns associated with different types of non-canonical junctions. For PEDV, SARS-CoV-2, and MERS-CoV, short-range junctions exhibit pronounced anti-diagonal enrichments in APA heatmaps, indicating multiple RNA-RNA interactions with convergent genomic directions. In contrast, this pattern is less discernible in the APA heatmap of PDCoV (Fig. 4A and Fig. S10A-C).

**Figure 4.**
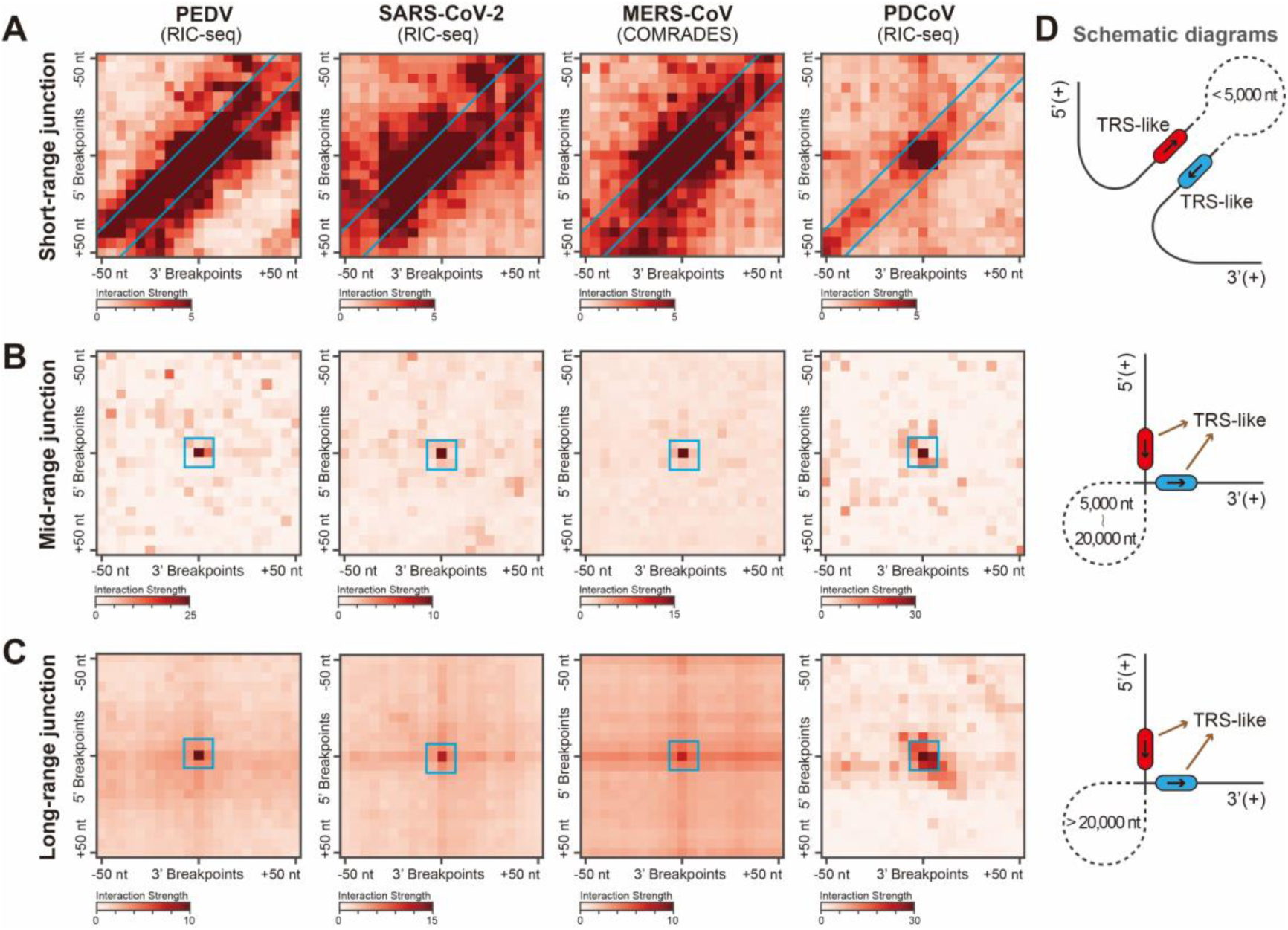
RNA architecture features of different types of non-canonical junctions. (A-C) APA plot shows the aggregate RNA-RNA interactions from short-range (A), mid-range (B) and long-range (C) junction site-flanking regions of PEDV, SARS-CoV-2, MERS-CoV and PDCoV. RNA-RNA interactions for PEDV, SARS-CoV-2 and PDCoV are detected by RIC-seq, and RNA-RNA interactions for MERS-CoV are detected by COMRADES. Highlighted points in the heatmap indicate genomic RNA-RNA interactions, while the anti-diagonal lines signify multiple interactions with convergent directions. (D) Schematic diagrams of RNA architecture underlying non-canonical junction: stem-loops contribute to the formation of short-range junctions; mid-range and long-range junctions are facilitated by distant genomic interactions.

As depicted in the APA heatmaps, convergent interactions indicate reverse complementary pairing between the flanking regions of short-range junction sites (Fig. 4A), contrasting with the parallel complementary base pairings observed between the flanking regions of TRS-L and TRS-B (Fig. 3B). The interaction strength between short-range junction site-flanking regions is significantly greater than that of controls across these coronavirus genomes (Fig. S10D and Fig. S11). Further examination reveals that some short-range TRS-like sequence pairs are located within the stem-loops, formed by reverse complementary base pairing (Fig. S12). Specifically, from an RNA-seq sample of Vero cells infected with SARS-CoV-2 (*2*), 5,014 non-canonical junctions were identified between three pairs of TRS-like sequences that are in spatial proximity due to a stem-loop spanning the genomic interval of 29,262-29,870 nt (Fig. S12A). These findings indicate that the co-localization of TRS-like sequence pairs with stem-loops could be a crucial factor in promoting short-range non-canonical junctions.

### Significant genomic interactions underlie mid- and long-range non-canonical junctions

For mid-range non-canonical junctions, a prominent central point is observed in the APA heatmaps for PEDV, SARS-CoV-2, MERS-CoV, and PDCoV (Fig. 4B and Fig. S10). Similarly, the APA heatmap patterns for long-range junctions in these coronaviruses resemble the mid-range interaction patterns, with PDCoV exhibiting an additional weak parallel diagonal line (Fig. 4C). Additionally, the average interaction strength between pairs of mid- or long-junction flanking regions significantly exceeds that of controls among these coronaviruses (Fig. S11B), highlighting the spatial proximity between pairs of TRS-like sequences. The closer evolutionary relationship between α- and β-coronaviruses, compared to δ-coronaviruses (*17*), may account for the slight structural differences in non-canonical junctions between PDCoV and the other viruses. Nevertheless, the underlying structural patterns for these three types of non-canonical junctions are broadly conserved across the coronaviruses (Fig. 4).

### Short-range non-canonical junctions benefit genomic deletion generation

Through multiple genome alignment, 105, 31, 17, and 8 genomic deletions (greater than 20 nucleotides missing) have been identified in the genomes of SARS-CoV-2, PEDV, MERS-CoV, and PDCoV, respectively (Table S2). Notably, the APA heatmaps for these genomic deletion regions show an obvious anti-diagonal enrichment for PEDV and PDCoV, and a weaker anti-diagonal enrichment for SARS-CoV-2 and MERS-CoV (Fig. 5A-D). The interaction strength between the opposing ends of the deletion regions is significantly higher than that of controls (Fig. S13), indicating an association between genomic deletions and stem-loops. Further examination reveals that 29.52%, 45.16%, and 62.50% of the deletion regions in SARS-CoV-2, PEDV, and PDCoV, respectively, overlap with stem-loops (Table S2).

**Figure 5.**
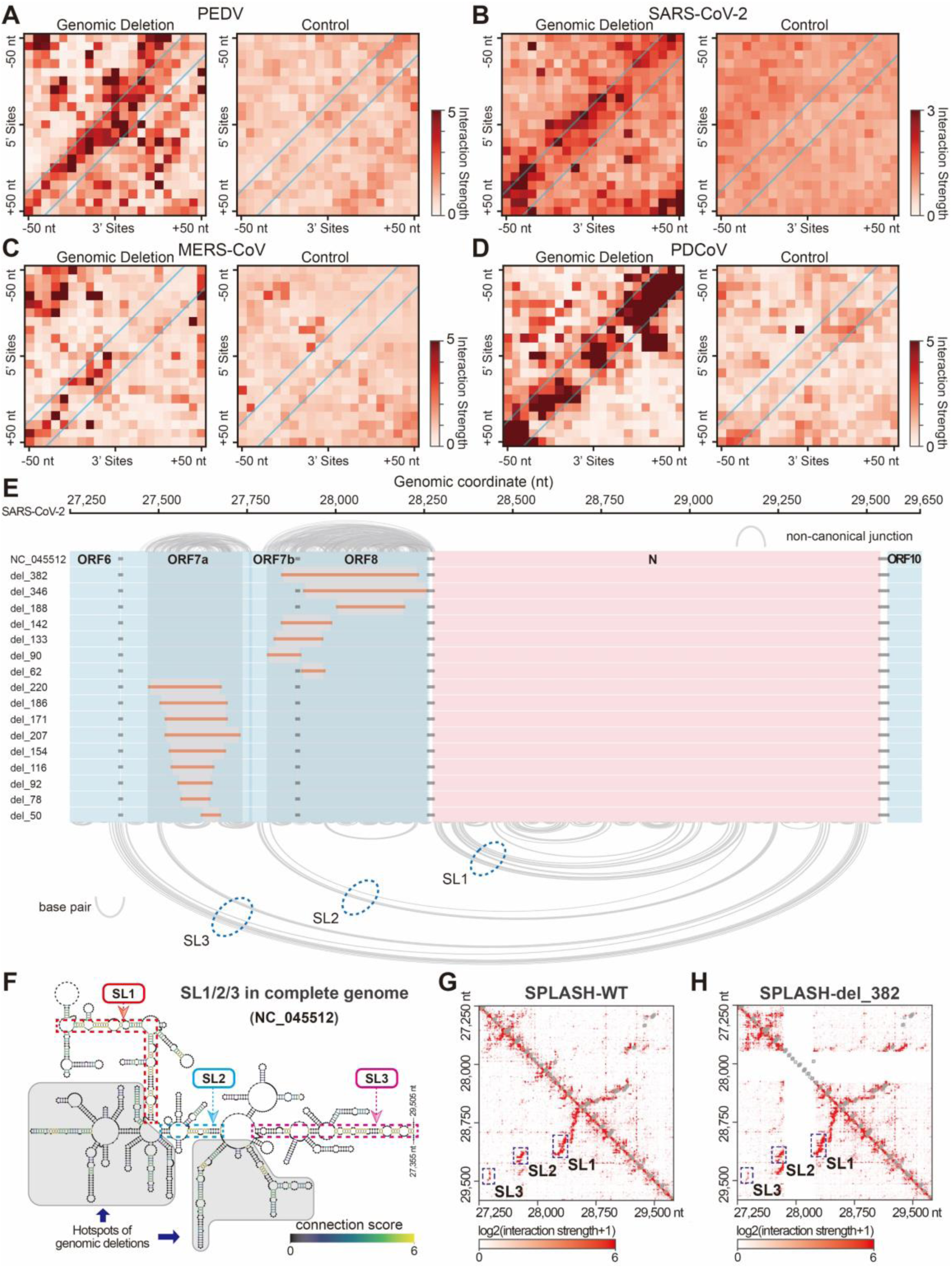
Genomic deletions overlap with stem-loops. (A-D) APA plot shows the aggregate RNA-RNA interactions from deletion flanking regions of PEDV (A), SARS-CoV-2 (B), MERS-CoV (C) and PDCoV (D). RNA-RNA interactions for PEDV, SARS-CoV-2 and PDCoV are detected by RIC-seq, and RNA-RNA interactions for MERS-CoV are detected by COMRADES. The random pairs of genomic sites spanning the same length as the genomic deletion regions were employed as the control. (E) The SL1/2/3 region in the SARS-CoV-2 genome is notable for containing 16 genomic deletions and featuring abundant non-canonical junctions. These shaded regions represent hotspots where genomic deletions frequently occur. (F) The RNA secondary structure of SL1/2/3 in the SARS-CoV-2 genome shows hotspots of genomic deletions, which are shaded. (G-H) Heatmaps depict the RNA-RNA interactions within the SL1/2/3 region of the SARS-CoV-2 genome, comparing interactions in the complete genome (G) with those in the defective genome (H). These genomic interactions are captured by SPLASH from Vero-E6 cells infected with different SARS-CoV-2 strains. The grey boxes at the top right of the heatmap represent base pairing in the RNA secondary structure.

An overlap between short-range non-canonical junction sites and specific stem-loop structures is also observed (Fig. 4A), suggesting that non-canonical junctions may play a role in initiating genomic deletions. We next examined the relationship between the locations of non-canonical junction sites and the regions where genomic deletions occur. In the genomes of PEDV, SARS-CoV-2, and MERS-CoV, the genomic distance between junction sites and deletion regions is significantly closer than in control regions (Fig. S14). For example, non-canonical junction sites are enriched within three representative stem-loops (SL1/2/3: 28,276 - 29,502 nt) in SARS-CoV-2 genome (Fig. S15). Notably, the SL1/2/3 region is also enriched with genomic deletions (16 deletions within 2,250 nt of SL1/2/3 region vs. 105 deletions within 29,903 nt of whole genome; *p*-value = 9.23e-93, Chi-square test) in the accessory protein genes (ORF7a, ORF7b and ORF8) of SARS-CoV-2 (Fig. 5EF and Table S3). Similarly, a stem-loop (3,260 - 3,450 nt) in the PEDV genome (MK584552.1), overlapping with 10 genomic deletions of ORF1a, is enriched with non-canonical junctions (Fig. S16 and Table S4). These findings suggest that generation of genomic deletions could be driven by short-range non-canonical junctions.

### Role of SL1/2/3 on ORF10 gene transcription in SARS-CoV-2 genome

The sequences of SL1/2/3 are highly conserved across different SARS-CoV-2-related genomes, with a sequence identity ranging from 0.84 to 0.93 (Fig. S17). Predictions of complementary base pairings indicate that the SARS-CoV-2-related SL1s share similar structures and the same number of base pairs (Fig. S18A). Additionally, the entire SL1/2/3 can be separately identified in the genome-wide interactions of SARS-CoV-2, including the strain with complete genome (hCoV-19/Singapore/2/2020) and the strain with ORF8 deletion (hCoV-19/Singapore/12/2020), as captured by SPLASH (Fig. 5FG). These data reveal the stability of SL1/2/3 across various SARS-CoV-2-related genomes.

In SARS-CoV-2 genome, ORF10 is the gene closest to 3’UTR and lacks TRS-B. Coincidentally, the 5’ parts of SL1/2/3 are located downstream of the TRS-B of genes such as N, ORF7a, and ORF7b, and its 3’ parts are located upstream of ORF10 (Fig. 5E). This arrangement suggests that the TRS-B sites of other genes could be transferred to ORF10 through SL1/2/3-mediated non-canonical junctions (Fig. 6A). This hypothesis is supported by the identification of sgRNAs, containing ORF10, TRS-B of other genes and leader sequence, from nanopore sequencing data of SARS-CoV-2-infected Vero cells (*2*) (Fig. 6B and Fig. S18B). Furthermore, we developed the Corona-mRNA algorithm to calculate Transcripts Per Million (TPM) using nanopore sequencing reads to assess the transcription level of ORF10 gene in SARS-CoV-2 (Fig. 6A and Fig. S19AB). An average of 11.37% ORF10 sgRNAs arise through double junctions among all nanopore samples (Fig. S19C). These findings suggest that the stable SL1/2/3 might act as a structural regulatory element for ORF10 transcription.

**Figure 6.**
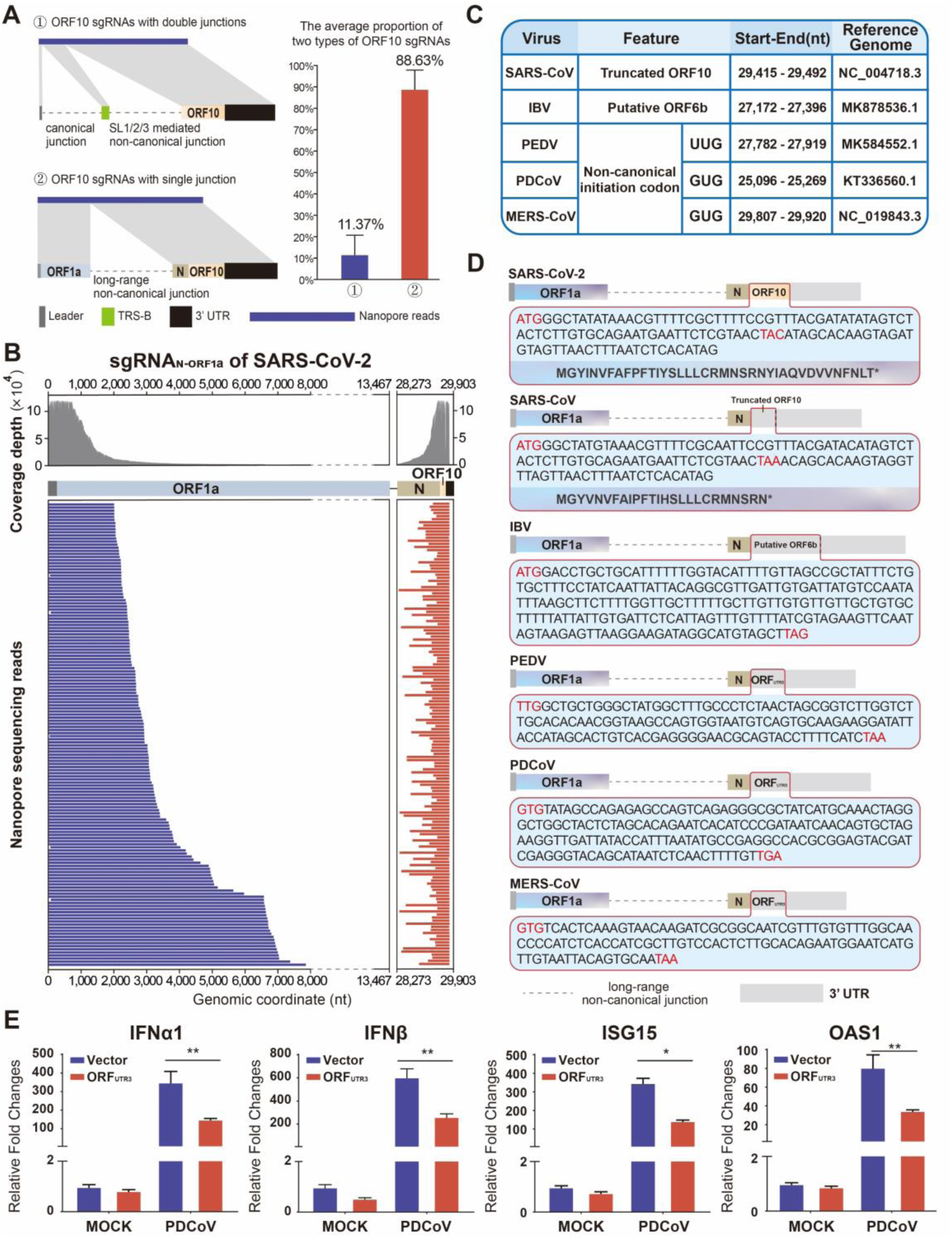
Conserved long-range non-canonical sgRNAs across coronaviruses with the similar function of immune responses. (A) Two types of non-canonical ORF10 sgRNAs detected from nanopore sequencing reads of SARS-CoV-2 infected Vero cells. One is formed by single junction between N gene and ORF1a, and the other by double junctions including the canonical junction and SL1/2/3-mediated non-canonical junction (left panel). The average proportion of ORF10 sgRNAs with single junction and double junctions in all nanopore samples, respectively (right panel). (B) The sgRNA_N-ORF1a_ of SARS-CoV-2 detected by nanopore sequencing reads. (C) Characterization of potential gene identified in the 3’ UTR of different coronavirus genomes. The initiation codon GUG or UUG with a low translation efficiency is termed as non-canonical initiation codon here. (D) Illustrative representations of sgRNA_N-ORF1a_ in the genomes of different coronaviruses. And the detailed nucleotide sequence of each potential gene is given. (E) LLC-PK1 cells were transfected with the ORF_UTR3_ plasmid or empty vector and were then infected with PDCoV. The transcription levels IFN-α1, IFN-β, ISG15 and OAS1 were monitored by qRT–PCR.

### Non-canonical sgRNAs with single junction between N gene and ORF1a are conserved across different coronaviruses

Besides the generation through double junctions, the majority of ORF10 sgRNAs are formed by single junction between the N gene and ORF1a among all SARS-CoV-2 samples (Fig. 6A and Fig. S19C). These sgRNAs, here designated as sgRNA_N-ORF1a_, are distinctly characterized by this unique junction (Fig. 6B). Notably, conserved junctions between N gene and ORF1a are also observed in other coronaviruses (Fig. S20A), leading to the formation of sgRNA_N-ORF1a_ with a 3’ UTR rather than an ORF with a defined function. Furthermore, conserved interactions between these two genomic regions were detected across different coronaviruses (Fig. S20B), implying that these interactions might contribute to the formation of sgRNA_N-ORF1a_.

We further investigated the correlation between the transcription level of sgRNA_N-ORF1a_ and RNA-RNA interaction strength using nanopore sequencing data (*3*) and simplified-SPLASH data (*13*) from SARS-CoV-2 at 24 hpi and 48 hpi (Figs. S20 and S21). A 12.76% decrease is observed in the genomic interaction strength between N gene and ORF1a at 48 hpi compared to 24 hpi, which corresponds to a significant reduction in the TPM of sgRNA_N-ORF1a_ (Fig. S22AB). In contrast, in negative samples captured by simplified-SPLASH without using RNA ligase, the strength of these interactions increases by 14.68% at 48 hpi compared to 24 hpi (Fig. S22C), indicating an actual decrease in interactions between N gene and ORF1a at 48 hpi. These findings indicate that the transcription level of sgRNA_N-ORF1a_ coincides with alterations in RNA-RNA interaction strength.

### Evolving genes with similar function are identified within sgRNA_N-ORF1a_ of different coronaviruses

Emerging evidence supports the coding ability of ORF10 within sgRNA_N-ORF1a_ of SARS-CoV-2, and ORF10 protein has been identified with the function of suppressing the antiviral innate immune response (*18–21*). A truncated version of ORF10, denoted as ORF10_truncated_, is found in the sgRNA_N-ORF1a_ of SARS-CoV with the termination codon (TAA) at 76 nt (Fig. 6CD). ORF10_truncated_, which shares homology with the 117 nt ORF10 of SARS-CoV-2 (Fig. 6D), is located in the 3’ UTR region; however, its function has not been reported. The mutation to TAC in the ORF10 of SARS-CoV-2 is considered to be crucial for encoding the functional protein (*22*, *23*). These findings indicate that the evolving gene within the 3’ UTR could potentially gain coding ability through base mutations.

Except for a putative ORF6b with undefined function in the 3’ UTR of IBV genome, no known open reading frames (ORFs) were identified in the 3’ UTR of PEDV, PDCoV, and MERS-CoV genomes (Fig. 6C). The 3’ UTR lengths of these coronaviruses are all longer than that of SARS-CoV-2 (Fig. S23A). Particularly, the 3’ UTR within the PDCoV genome is 383 nt long, surpassing the combined length of the 3’ UTR (155 nt) and ORF10 (117 nt) within the SARS-CoV-2 genome. This suggests that a short gene might exist within these relatively long 3’ UTRs. Potential ORFs with non-canonical initiation codons (UUG or GUG) were predicted in the 3’ UTRs of MERS-CoV, PEDV, and PDCoV genomes, which could acquire the canonical initiation codon (AUG) through a single base mutation (Fig. 6CD). These potential ORFs within the 3’ UTR are termed ORF_UTR3_ here.

To investigate the role of ORF_UTR3_ in the antiviral innate immune response, we engineered a variant ORF_UTR3_ of PDCoV by mutating the non-canonical initiation codon GUG with the canonical AUG. Subsequently, LLC-PK1 cells were transfected with this modified ORF_UTR3_ construct (pCAGGS-ORF_UTR3_) alongside a control vector (pCAGGS-Flag) and whole-cell lysates were assessed using quantitative real-time PCR (qRT–PCR). The overexpression of ORF_UTR3_ significantly reduced the transcription of IFN-α1 and IFN-β upon PDCoV infection (Fig. 6E). Furthermore, the mRNA levels of ISG15 and OAS1 in ORF_UTR3_-overexpressing cells are lower compared to the control in the presence of PDCoV (Fig. 6E). An increase in the quantity of N sgRNAs, indicating enhanced viral replication, is also observed in LLC-PK1 cells overexpressing ORF_UTR3_ (Fig. S23B). The inhibitory effect of ORF_UTR3_ on the antiviral innate immune response is consistent with the reported function of ORF10 in SARS-CoV-2 (*20*).

## Discussion

Coronavirus produces individual canonical sgRNA encoding structural and accessory proteins through discontinuous transcription. This process benefits virus proliferation by precisely controlling the production ratio of these proteins to meet the virus’s specific requirements. In this study, we found that canonical sgRNAs are not consistently predominant, and the prevalence of non-canonical sgRNAs varies depending on the infection phase and the specific viral strain (Fig. 2A and Fig. S2CD). The roles of non-canonical transcripts are noted in several studies, but understanding how the virus controls their production relative to canonical sgRNAs is a tough problem. APA analysis reveals an antiparallel arrangement for the RNA-RNA interaction between TRS-like sequences pairs of short-range non-canonical junctions. In contrast, RNA-RNA interactions with same genomic orientation are observed surrounding the TRS pairs for canonical transcriptional junctions. Despite the fundamental evolutionary principle of antiparallel orientation in nature, parallel stretches exist and they can be found both in vivo and in vitro (*24*). The existence of parallel-stranded DNA duplexes was first confirmed using Raman spectroscopy and chemical methylation (*25*), and subsequently refined by infrared and NMR spectroscopy (*26–28*). Parallel DNA or RNA duplexes in various genomes, including *Drosophila* and *Escherichia coli*, are involved in biological functions such as mRNA processing and gene silencing, with parallel RNA strategies proving more effective than antisense RNA for gene targeting (*29–31*). We speculate that parallel RNA-RNA interactions may facilitate template switching between TRS-L and TRS-B more effectively; however, how the two RNA segments engage in parallel spatial interactions requires further investigation.

Deletions are common during the evolution of coronavirus, and previous studies suggest that deletions in SARS-CoV-2 are assumed to arise at template-switching hotspots (*32*, *33*). We investigated the RNA-RNA interaction patterns in regions where deletions exceed 20 nucleotides in length to uncover the precise RNA architectures that drive genomic deletions in the virus. In the SARS-CoV-2 genome, the SL1/2/3 region, which is enriched with 16 genomic deletions, overlaps with stem-loops mediating abundant non-canonical junctions (Fig. 5E). Similarly, in PEDV genome, a region containing 10 genomic deletions in ORF1a overlaps with a stem-loop and is also enriched with non-canonical junctions (Fig. S16). Basing on these findings, we propose that short-range non-canonical junctions mediated by stem-loops may be one of the driving factors for long genomic deletions. Advances in high-resolution RNA architecture detection may enable the exploration of finer details, including the driving force for deletions shorter than 20 nucleotides.

Genomic deletions could be an important way to drive viral evolution and develop attenuated vaccines. ORF8 is a hypervariable gene rapidly evolving among SARS-related coronaviruses, with a tendency to undergo deletions (*34*). Notably, infections with SARS-CoV-2 strains harboring ORF8 deletions are associated with milder clinical symptoms and less systemic release of pro-inflammatory cytokines (*35*). Virus attenuation based on passaging culture is a traditional strategy for vaccine development (*36*, *37*). The caPEDV (JQ023162; isolated from a commercial vaccine of GreenCross, South Korea), which contains a deletion within ORF3 gene, was obtained by passaging culture and showed an attenuated phenotype in pigs (*38*). Understanding the mechanisms of genomic deletion generation is beneficial for further predicting virus evolution and designing more stable attenuated vaccines.

Insertions, deletions and mutations are the main pathways for viruses to generate new genes and contribute to the virus’ adaptive evolution (*39*, *40*). The generation of conserved non-canonical sgRNAs via junctions between N gene and ORF1a implies a possible mechanism of convergent evolution or co-evolution across different coronaviruses. This study elucidates the transcriptional mechanism of the ORF10 gene in the SARS-CoV-2 genome, highlighting its utilization of non-canonical junctions. Putative or evolving ORFs were identified within sgRNA_N-ORF1a_ of other coronavirus, and these evolving ORFs have the potential to encode novel proteins through single-nucleotide mutations. Despite our observation that overexpression of ORF_UTR3_ effectively attenuated the innate immune response upon PDCoV infection (Fig. 6E), additional experiments in viral replication assays are required to validate the influences of ORF_UTR3_ on viral virulence and pathogenicity. Whether these ORF_UTR3_ will persist in the process of viral evolution depends on the ongoing selection pressures between the virus and its host. Further studies are needed to determine whether a similar phenomenon occurs in other members of the Nidovirales order that utilize discontinuous transcription mechanisms.

## Materials and Methods

### Cell culture and virus infection

Vero E6 cells were maintained in Dulbecco’s modified Eagle’s medium (DMEM; Invitrogen, CA, USA), supplemented with 10% fetal bovine serum (FBS) and 1% penicillin streptomycin (Invitrogen, CA, USA). LLC-PK1 cells were cultured in Minimun Essential Medium (MEM; Cytiva), supplemented with 10% fetal bovine serum (FBS) and 1% penicillin streptomycin (Invitrogen, CA, USA). Cells was cultured at 37℃ in an atmosphere of 5% CO_2_ and 95–99% humidity. Vero E6 cells were infected with PEDV AJ1102 (GenBank accession: MK584552) at a MOI of 1.0 for 12 hours, while LLC-PK1 cells were infected to PDCoV CHN-HN-2014 (GenBank accession: KT336560) at an MOI of 1.0 for 10 hours. Infections were performed in DMEM (for PEDV) or MEM (for PDCoV) supplemented with 7.5 μg/mL trypsinization.

### RIC-seq library preparation

In the late stage of replication, infected cells were crosslinked with 1% formaldehyde for 20 minutes at room temperature and then quenched by the addition of glycine. The crosslinked cells were subsequently scraped off the plate and collected by centrifugation at 3,000 rpm for 10 minutes at 4°C. The subsequent library preparation was conducted as previously described (*41*), followed by sequencing on an Illumina HiSeq X Ten platform.

### RNA-seq library preparation

In the late stage of replication, infected cells were harvested. Total RNA was extracted using TRIzol (Invitrogen, USA) according to the manufacturer’s instructions. RNA-seq libraries were constructed from 2 μg of RNA using the VAHTS Universal V6 RNA-seq Library Prep Kit (Vazyme, CHN), and subsequently sequenced on the MGISEQ-2000 platform with paired-end 150 bp reads.

### Plasmid constructs

Total RNA was extracted from PDCoV-infected LLC-PK1 cells. The ORF_UTR3_ sequence was then mutated to convert non-canonical start codons (UUGs) to canonical start codons (AUGs) using targeted mutation primers (forward primer: CCGGGTACCatgtatagccagagagcc; reverse primer: CCGCTCGAGtcaacaaaagttgagatta). After mutation, the mutated ORF_UTR3_ was cloned into the pCAGGS-Flag vector. All enzymes used for the cloning procedures were purchased from Vazyme, Nanjing, China.

### Overexpression of ORF_UTR3_ and qRT-PCR

The pCAGGS-ORF_UTR3_ and pCAGGS-Flag were transfected into LLC-PK1 cells, respectively, and the cells were infected with 0.1 MOI of PDCoV 24 hours later. Total intracellular RNA was extracted at whole-cell lysates, and mRNA was quantified as described previously (*20*). And qRT-PCR was used to detect mRNAs of IFN-I genes, IFN-stimulated genes and viral N gene. The β-actin gene was used as an internal reference to ensure relatively accurate mRNA expression and to avoid system and random errors during sample processing. The primer sequences used are listed as follows:

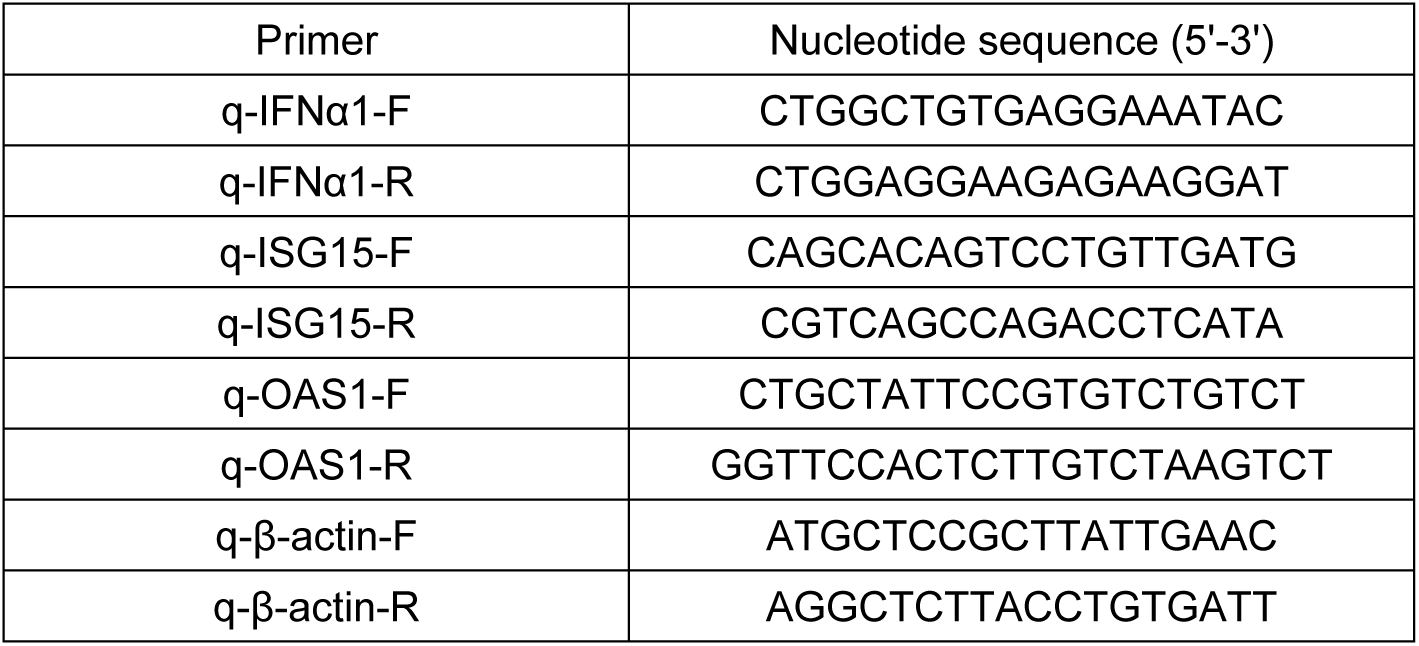

### Processing of RNA-RNA interactome and RNA-seq data

The pipeline developed for RIC-seq data analysis (*41*) was employed to uniformly process experimental data from RNA-seq, RiC-seq, COMRADES, SPLASH, and simplified-SPLASH, which essentially count the number of chimeric RNA reads formed by ligase proximity ligation or presented in sgRNAs. Ultimately, the outcomes of genome-wide RNA-RNA interactions and junction sites within sgRNAs were stored in ".hic" and ".mcool" formats. We refer to adopt the pipeline (*41*) of analyzing RIC-seq data for unified processing, which is divided into the following 4 steps:

1. Fastp (*42*) was used for quality control of the sequencing data, and then software such as STAR (*43*) and bwa (*44*) were used to identify reads that were aligned to the reference genome;
2. Paired reads were identified the from the aligned reads that includes STAR and BWA outputs;
3. Considering the existence of intra-RNA molecule and RNA-RNA molecule interactions in the paired reads, the process is different for different types of data. If it is RNA-RNA interactome data, then the paired reads are categorized as normal and chimeric, where intra-RNA molecule interactions are regarded as normal and RNA-RNA intermolecular interactions are regarded as chimeric. If it is RNA-seq data, then this step obtains information about the junction sites of discontinuous transcripts;
4. After identifying the junction sites of chimeric reads, an RNA-RNA interaction matrix in ".hic" format can be constructed, while the Knight-Ruiz algorithm is used to balance this interaction matrix to eliminate experimental errors along the viral genome (*14*);
5. Pseudo-topological associating domains (pTADs) were divided based on the RNA-RNA interactions matrix. Within each pTAD, RNA secondary structure was predicted using RNAstructure (*45*) using RNA-RNA interactions as constraints.

RNA-seq data are mainly used to analyze the sgRNAs of coronaviruses genome, which is essentially also a search for chimeric reads, and thus can be processed using the same pipeline for RNA-RNA interactome data. It is worth noting that the chimeric reads identified in the third step of RNA-seq data analysis should be regarded as sgRNAs junctions formed by discontinuous transcription, and this part of the chimeric reads can be removed when analyzing the RNA-RNA interactome data, so as to more accurately reflect the real RNA-RNA interactions.

### Processing of nanopore-seq data and workflow of Corona-mRNA

For nanopore sequencing data, we developed a custom pipeline to quantify the transcribed mRNAs for each gene, leveraging the comprehensive sgRNA profiles identified through direct RNA sequencing approach. Nanopore reads are stored in the form of electrical signals in ".fast5" files, which can be converted to ".fastq" files with the option of NanoPlot and NanoFilt software (*46*) for data quality control. The reference genome was then indexed using minimap2 (*47*) software (parameter: -d), followed by aligning the data to the reference genome (parameter: -ax splice -c). For the ".sam" files aligned to the genome, the reads abundance and expression for each gene of coronavirus were calculated using Corona-mRNA.

Corona-mRNA utilizes the ".sam" file generated from sequence alignment as input and outputs the abundance of mRNAs or gene expression levels. As shown in Fig. S19A, Corona-mRNA is divided into four main steps:

1. CIGAR (Compact Idiosyncratic Gapped Alignment Report) information are extracted from the ".sam" file and converted into sgRNAs or gRNAs information;
2. Screen sgRNAs that contain both the leader sequence, a complete gene reading frame, and the 3’ UTR region. These sgRNAs formed by a single junction of TRS-L and TRS-B are canonical, the rest are non-canonical;
3. The sgRNAs with the potential to express the same protein are grouped into the same category, where the gene closest to the leader sequence in the sgRNAs is considered to be transcribed;
4. Calculate the abundance of mRNAs and TPM of each gene by canonical and non-canonical junctions, and then the results of the two conditions are combined.

### Identification of canonical and non-canonical junctions

We identified canonical and non-canonical junctions from RNA-seq data. For canonical junction sites, we screened for pairs of junction sites that TRS-L is involved in and the number of junctions exceeded 2,000 or 1/100 of the maximum junctions. For non-canonical junction sites, we screened for pairs of junction sites with junctions exceeding 10 and distances exceeding 100 nucleotides. For RNA-seq datasets in Fig. S4B, we classified non-canonical junctions using a uniform criterion, both using 5,000 nt and 20,000 nt as the dividing genomic distance.

### Sequence consistency

The transcription regulatory sequences TRS-L and TRS-B consist of approximately 6 conserved RNA bases (specifically ACGAAC in SARS-CoV-2), inducing a pause in RdRp synthesis of genomic RNA at the TRS-B region, leading to a template switch to TRS-L within the leader sequence. Consequently, an investigation was conducted to evaluate the sequence conservation of RNA pairs (6 nt before or after the junction site) involved in the formation of non-canonical junctions across coronavirus genomes.

Sequence consistency is employed to assess the similarity between pairs of positive-sense strand RNAs that have been identified to possess non-canonical junctions between them. The Bio.pairwise2 module (v1.81) in Python (v3.9) is used to evaluate the consistency between two RNA sequences, and "localms" is chosen as the "alignment type".

Since G is able to form base complementary pairing with U in RNA, C in 3’ RNA can match U in 5’ RNA, and A in 3’ RNA can match G in 5’ RNA. Therefore, we constructed a match matrix "M" for RNA bases (A, U, C and G) alignment as follows:

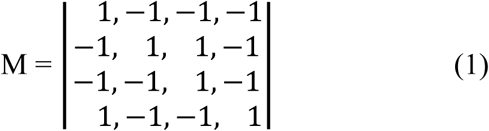

As shown in the matching matrix "M", 1 point is given for a match and 1 point is penalized for a mismatch. In addition, there is a 0.5 penalty point for either opening a gap or extending a gap.

### Aggregate peak analysis (APA)

We perform APA for canonical/non-canonical junctions and genomic deletions based on 5 nt resolution RNA-RNA interaction matrices. To measure the aggregate enrichment of a set of junction sites in an interaction matrix, the heatmap of sum of a series of submatrices derived from that interaction matrix are plotted. Each of these submatrices is a 100 nt × 100 nt square centered at a single junction site in the interaction matrix. The resulting APA heatmap shows the total number of interactions that lie within the entire set of junction sites at the center of the matrix. The APA heatmap of junction sites only include whose loci are at least 100 nt apart to avoid the effects of proximal RNA-RNA interactions. And random pairs of genome sites with equivalent numbers and lengths to junction sites were employed as the control.

Structural analysis of the genomic regions encompassing deletions was conducted using APA, focusing on RNA-RNA interactions within 50-nucleotide regions surrounding the deletion boundaries. Random pairs of genomic sites with equal length to the actual deletion regions were employed as the control. Most of the deletion sites in coronaviruses are less than 100 nt, so the "observed over expected" matrix, which eliminates the effect of genomic distance, is used to plot the APA heatmap. Points or lines highlighted in the APA heatmap, which indicate the underlying structural pattern between the two sites involved in the junctions or deletions.

### Prediction of complementary pairing between pairs of RNA sequences

RNAfold (web server: http://rna.tbi.univie.ac.at/cgi-bin/RNAWebSuite/RNAfold.cgi) was used to predict the complementary pairing between pairs of genomic RNAs. In predicting complementary pairing of pairs of genomic RNAs in the same direction (TRS-B and TRS-L flanking region), we reverse sequences in the 3’ parts of genomic RNA, while the pairs of genomic RNAs in convergent direction (SL1) is not required. Predictions were made with default parameters and "Folding Constraints" were added. Specifically, "<" was set as a constraint of each base in the 5’ part of RNA sequences, and ">" was set as a constraint of each base in the 3’ part of RNA sequences. Among the predicted results, we screen for complementary pairing that best matches the RNA-RNA interactions, which can finally be visualized using VARNA (*48*) (v3.9) or RNAstructure (v6.4).

### Identification of genomic deletions

A total of 833 PEDV genome sequences, 1,110 SARS-CoV-2 genome sequences, 657 MERS-CoV genome sequences and 213 PDCoV genome sequences were downloaded from NCBI and GISAID. Then, those genome sequences of each virus were aligned using MAFFT (v7.505). We defined MK584552, NC_004718, NC_019843 and KT336560 as reference genomes for PEDV, SARS-CoV-2, MERS-CoV and PDCoV, respectively. Finally, manually examined regions with greater than 20 nucleotides missing on other genomes relative to the reference genome are treated as genomic deletions.

### Prediction of potential ORFs in the 3’ UTR of coronaviruses

SnapGene software (www.snapgene.com) was used to predict potential ORFs in the 3’ UTRs of PEDV, MERS-CoV, IBV, and PDCoV. In order to predict ORFs within the 3’ UTR region, the minimum number of amino acids was set to 10, and GUG or UUG could be used as the initiation codon in addition to AUG. The predicted ORFs were then screened for their location within most of the long-range non-canonical sgRNAs. Finally, a manual examination is required to determine whether the predicted ORFs are reasonable.

## Acknowledgement

We thank Liheng Ding and Shuhua Yang for their support and the valuable suggestions they offered throughout the database construction process. We thank Professor Wen Su and Di Liu for editorial help and comments on the manuscript. We also thank Jiahui Guo for his advice on analysis and comments on the manuscript.

## Author contributions

Z.W., D.W. and L.L. conceived the project; S.X. provided virus samples of PEDV and PDCoV; D.L. and J.S. performed RNA-seq and RIC-seq experiments for PEDV and PDCoV; L.G. performed plasmid constructs and qRT-PCR for ORF_UTR3_ in PDCoV; ZW performed the computational analysis. Y.D.; Z.H., L.C., K.P. and Y.W. contribute to data collection and pre-processing; L.L., D.W. and F.W. advised on data analysis; ZW wrote the manuscript with input from all authors; D.W. and L.L. revised the manuscript.

## Data and materials availability

The RIC-seq and RNA-seq data for PEDV and PDCoV have been deposited in the National Genomics Data Center (https://ngdc.cncb.ac.cn/) under the BioProject accession number PRJCA026550 that are publicly available. The public data used in this study were collected from NCBI, EBI, BIG, and OSF, and the processed data were stored in CODE. CODE is available at "https://webofgroup.cn/dgwei/hngene". CODE collected RNA-RNA interactomes (COMRADES, SPLASH, simplified-SPLASH and RIC-seq) and transcriptomes (RNA-seq and Nanopore-seq) from 14 coronaviruses including HCoV-229E, HCoV-NL63, PEDV, TGEV, SADS-CoV, HCoV-HKU1, HCoV-OC43, MERS-CoV, MHV, PHEV, SARS-CoV, SARS-CoV-2, IBV and PDCoV.

## Code availability

Homemade scripts for analyzing junction sites within coronavirus sgRNAs and RNA architecture could be found at https://github.com/WenZi0809/RNA-architecture-underlies-discontinuous-transcription-and-evolution-of-coronavirus.

