## Supplemental information for "RNA Architecture Underlies Discontinuous Transcription and Evolution of Coronavirus"

Zi Wen *et al.*

### Supplementary Text

**Figure S1 – Figure S22: Captions are included below each of the supplementary figures.**

**TABLE S1. Proportion of non-canonical junctions at different genomic distances.**

**TABLE S2. Deletions obtained based on multiple sequence alignment.**

**TABLE S3. Locations of enriched genomic deletions in SARS-CoV-2 genome (NC_045512.2).**

**TABLE S4 Locations of enriched genomic deletions in PEDV genome (MK584552).**

**Data S1. Basic information of RNA-seq samples.**

**Data S2. Basic information of nanopore samples.**

**Data S3. Genomic deletions identified from coronavirus genomes of PEDV, SARS-CoV-2, MERS-CoV and PDCoV.**

**Data S4. TPM for two types of ORF10 sgRNAs in nanopore samples of SARS-CoV-2.**

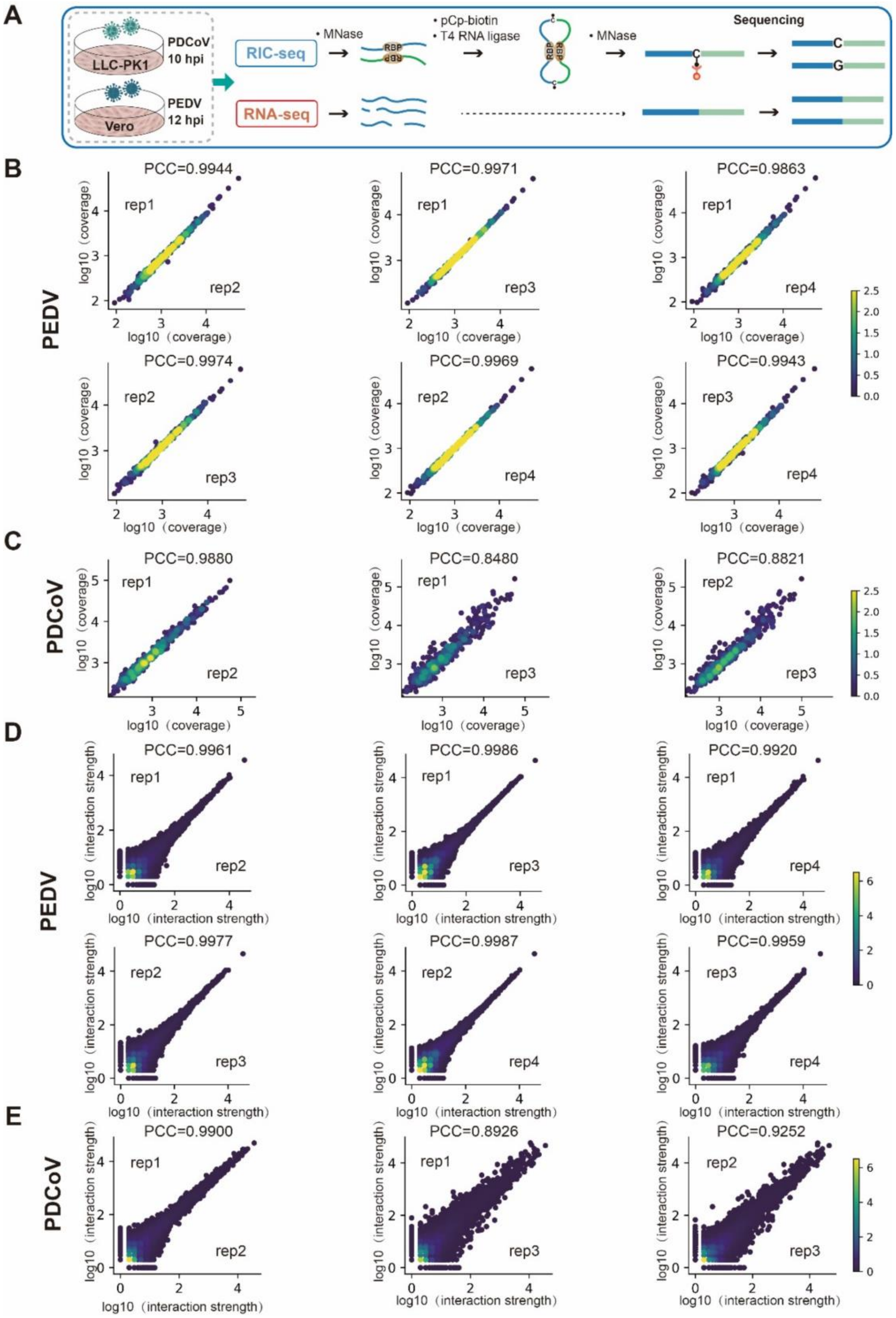


**Fig. S1 Key steps of the RIC-seq experiment and data quality.**

**A**, Schematic diagrams illustrate the key steps and experimental conditions for RNA-seq and RIC-seq experiments involving Porcine Epidemic Diarrhea Virus (PEDV) and Porcine Deltacoronavirus (PDCoV). **B-C**, Scatter plots revealed the correlation of chimeric read coverage along the PEDV (B) or PDCoV (C) genomes between each pair of biological replicates. **D-E**, Scatter plots revealed the correlation of interaction strength between each pair of biological replicates for PEDV (D) or PDCoV (E). The Pearson correlation coefficient (PCC) was calculated for each pair of replicates.

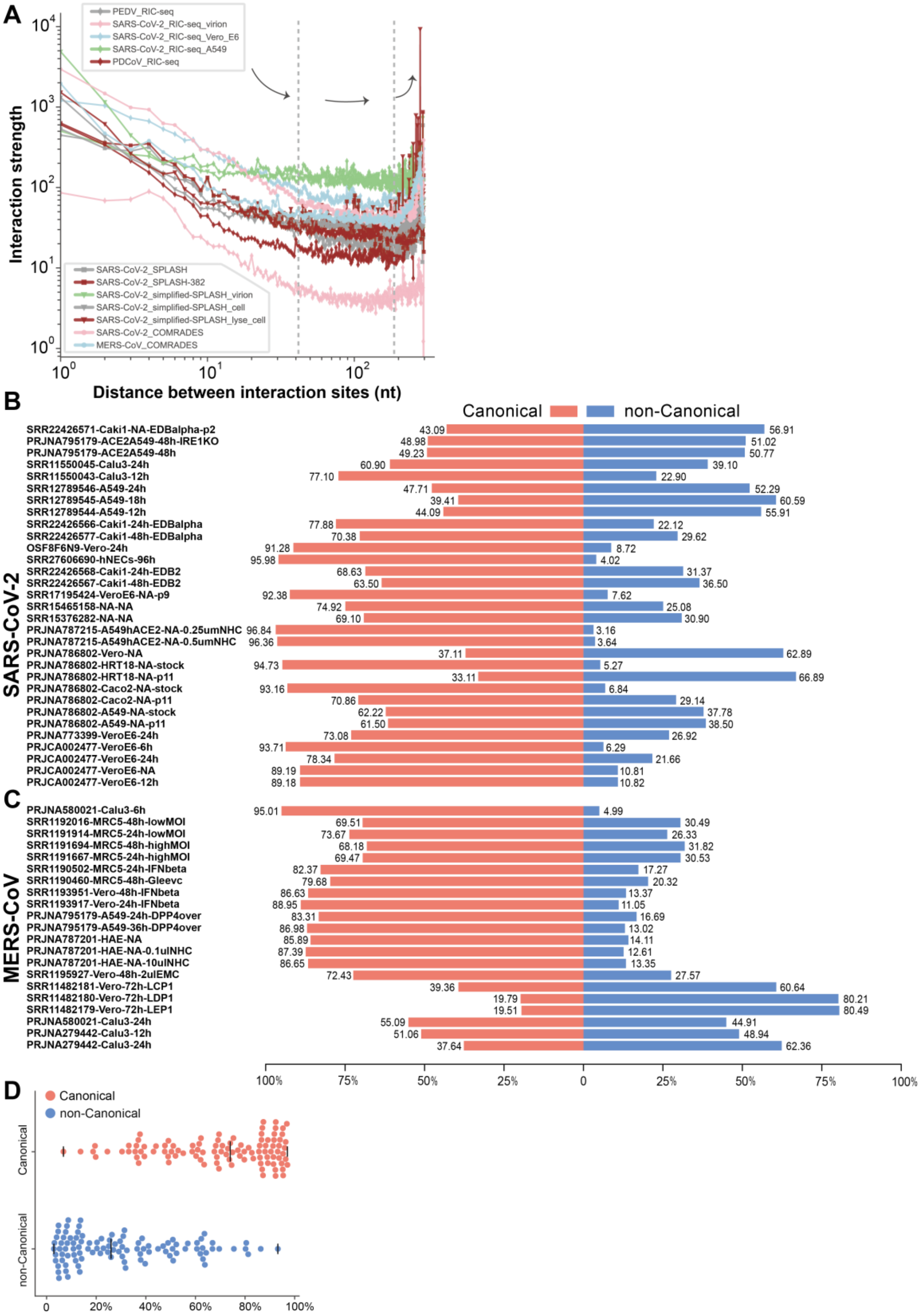


**Fig. S2 Statistics of RNA-RNA interaction data and transcriptome data.**

**A**, RNA-RNA interaction strength across a range of distances between pairwise interacting RNAs from different coronaviruses. **B-C**, Proportions of canonical and non-canonical junctions in RNA-seq samples of SARS-CoV-2 (B) and MERS-CoV (C). The SRA or BioProject accession number is added to the sample name. **D**, Distribution of proportions of non-canonical and canonical junctions based on 103 RNA-seq samples.

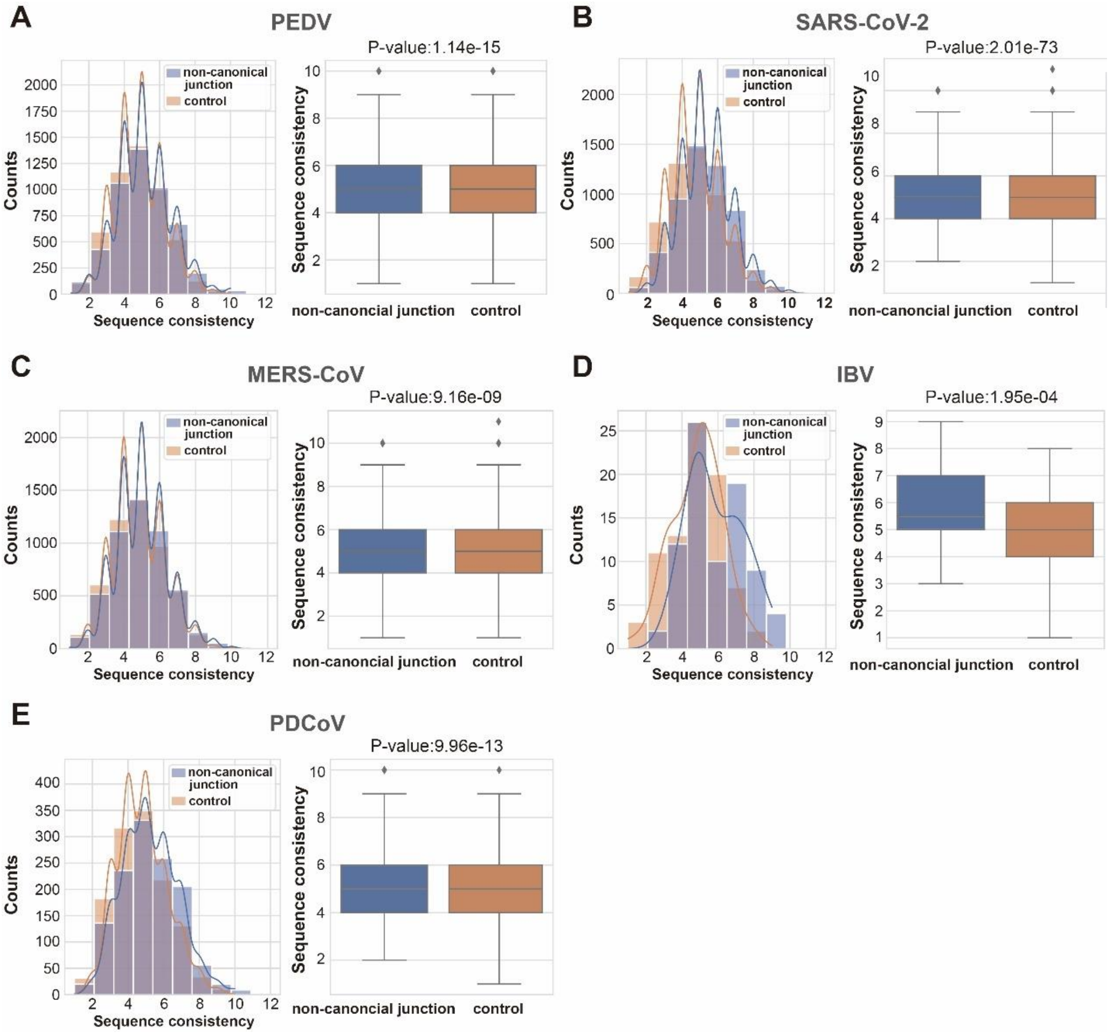


**Fig. S3 RNA sequence consistency of non-canonical junctions in different coronavirus genomes.**

**A-E**, Histogram and Boxplot of sequence consistency between pairs of genomic RNAs involved in non-canonical junctions in PEDV (A), SARS-CoV-2 (B), MERS-CoV (C), IBV (D) and PDCoV (E) genomes. The p-value of Wilcoxon rank-sum statistical test was calculated for each group.

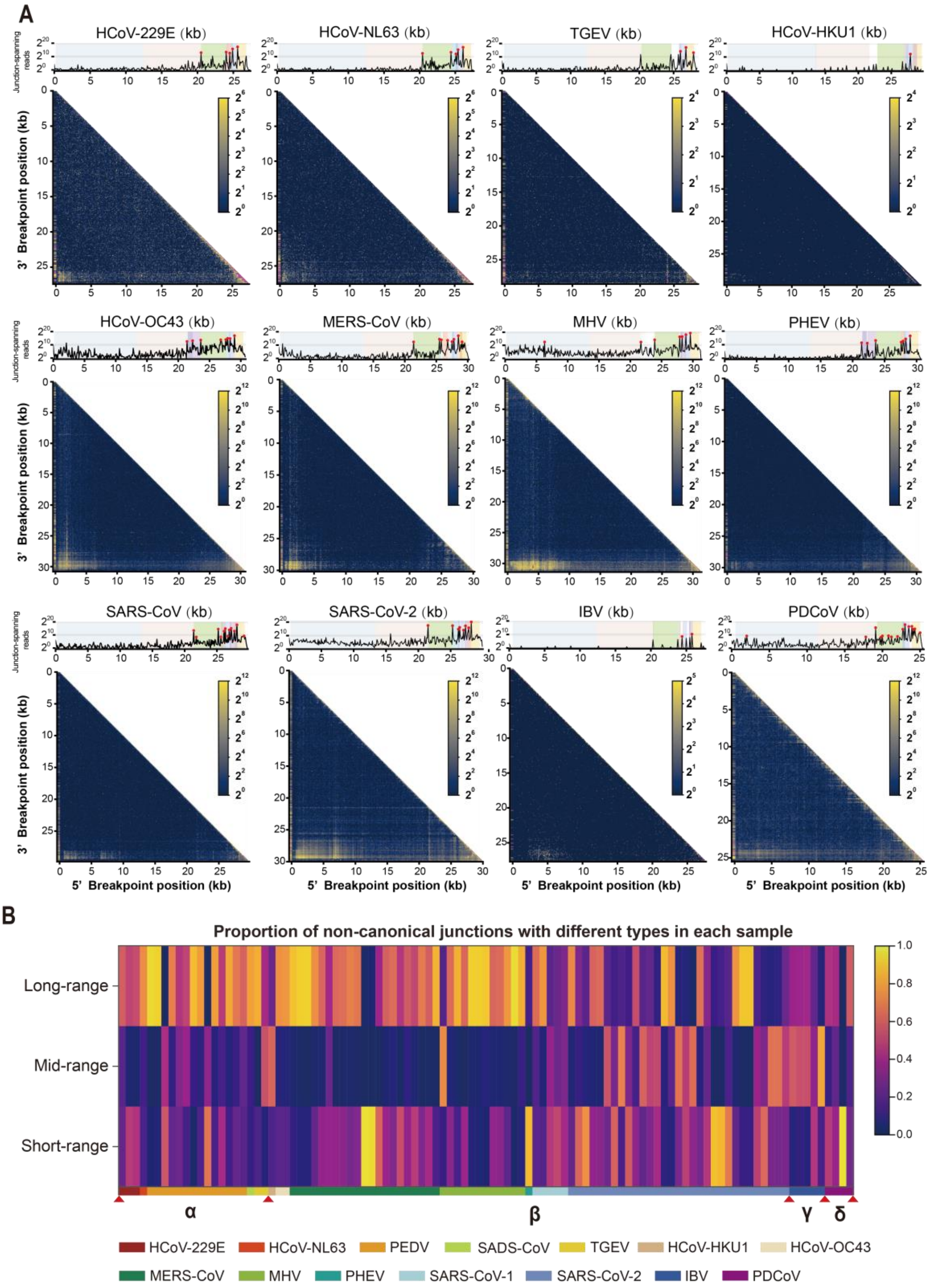


**Fig. S4 Distribution of junction sites in different coronavirus genomes.**

**A**, Heatmaps depicting the distribution of junction sites for different coronavirus genomes. **B**, Heatmap depicting the proportion of non-canonical junctions with different types among all RNA-seq samples.

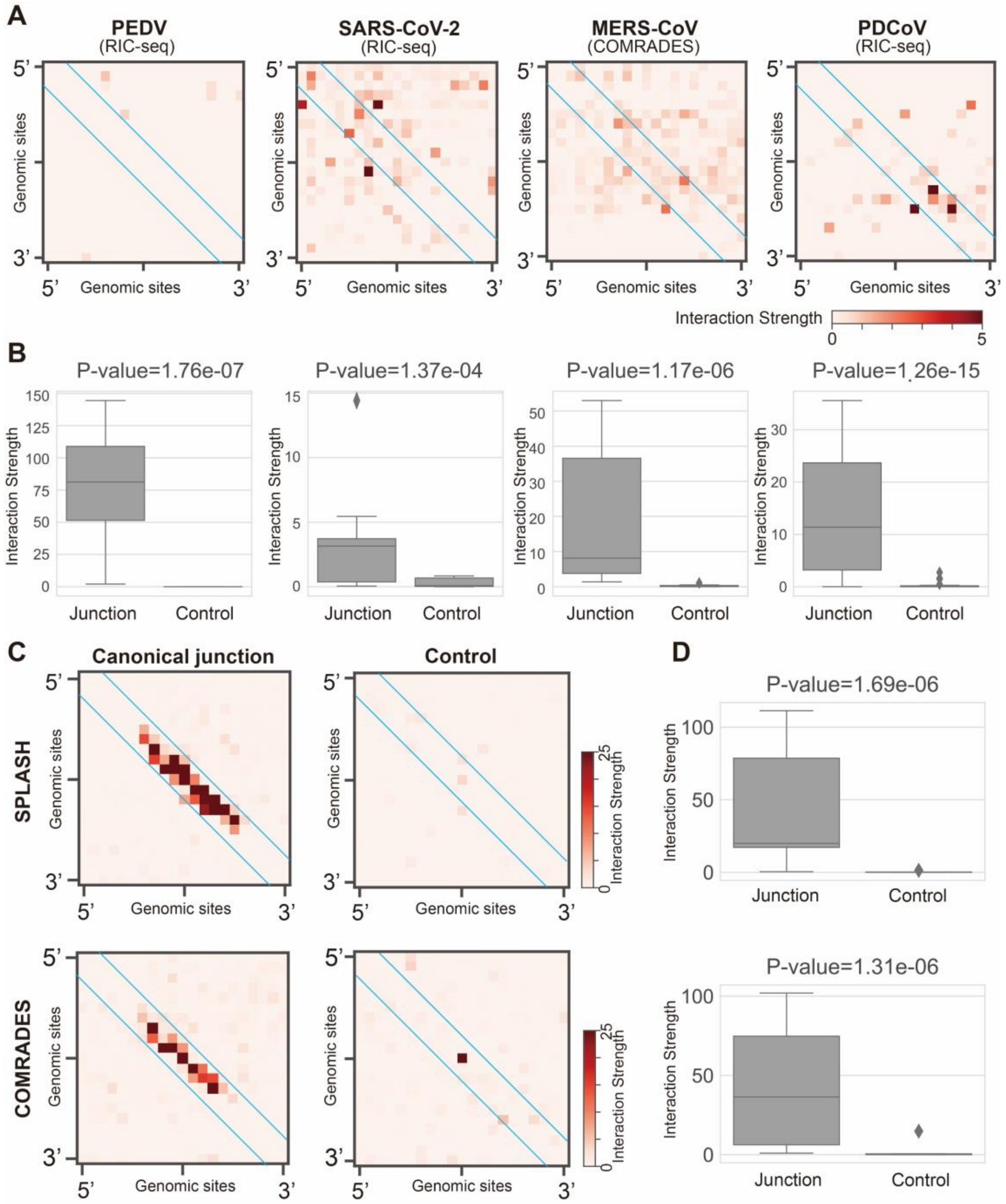


**Fig. S5 Aggregation peak analysis (APA) and comparison of interaction strength for canonical junctions.**

**A**, Random pairs of sequences spanning equivalent lengths as the canonical junctions are employed as the control. APA plot shows the aggregate RNA-RNA interactions from control groups. RNA-RNA interactions for PEDV, SARS-CoV-2 and PDCoV are detected by RIC-seq, and RNA-RNA interactions for MERS-CoV are detected by COMRADES. **B**, Boxplots depicting the RNA-RNA interaction strengths between canonical junction sites, compared with control groups. Interaction strengths were determined by selecting the regions delineated between the two blue lines in (A). **C**, APA plot shows the aggregate RNA-RNA interactions from canonical junction site-flanking regions and control groups of SARS-CoV-2. RNA-RNA interactions are captured by SPLASH and COMRADES, respectively. **D**, Boxplots depicting the RNA-RNA interaction strengths between canonical junction sites, compared with control groups. Interaction strengths were determined by selecting the regions delineated between the two blue lines in (C). The p-value of one-tailed t-test was calculated for each group.

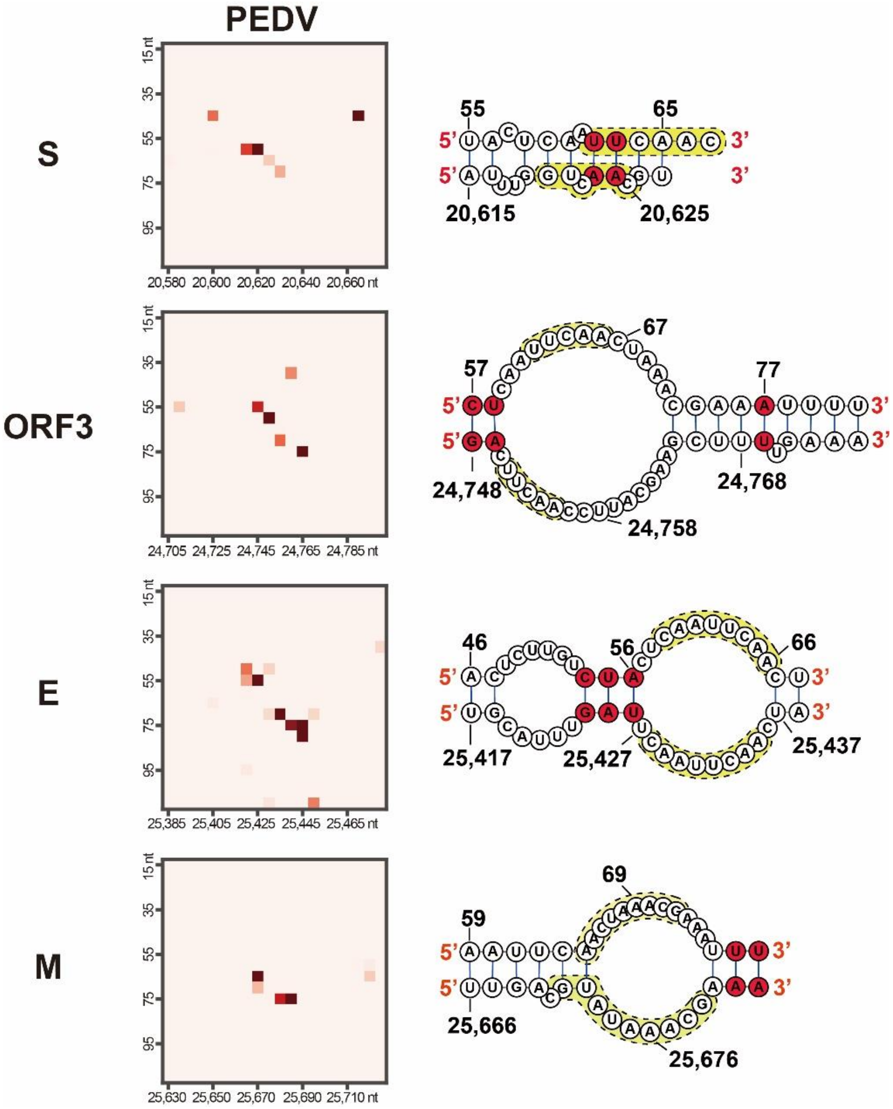


**Fig. S6 RNA-RNA interactions and base pairing between TRS-L flanking region and TRS-B flanking region for each gene in PEDV.**

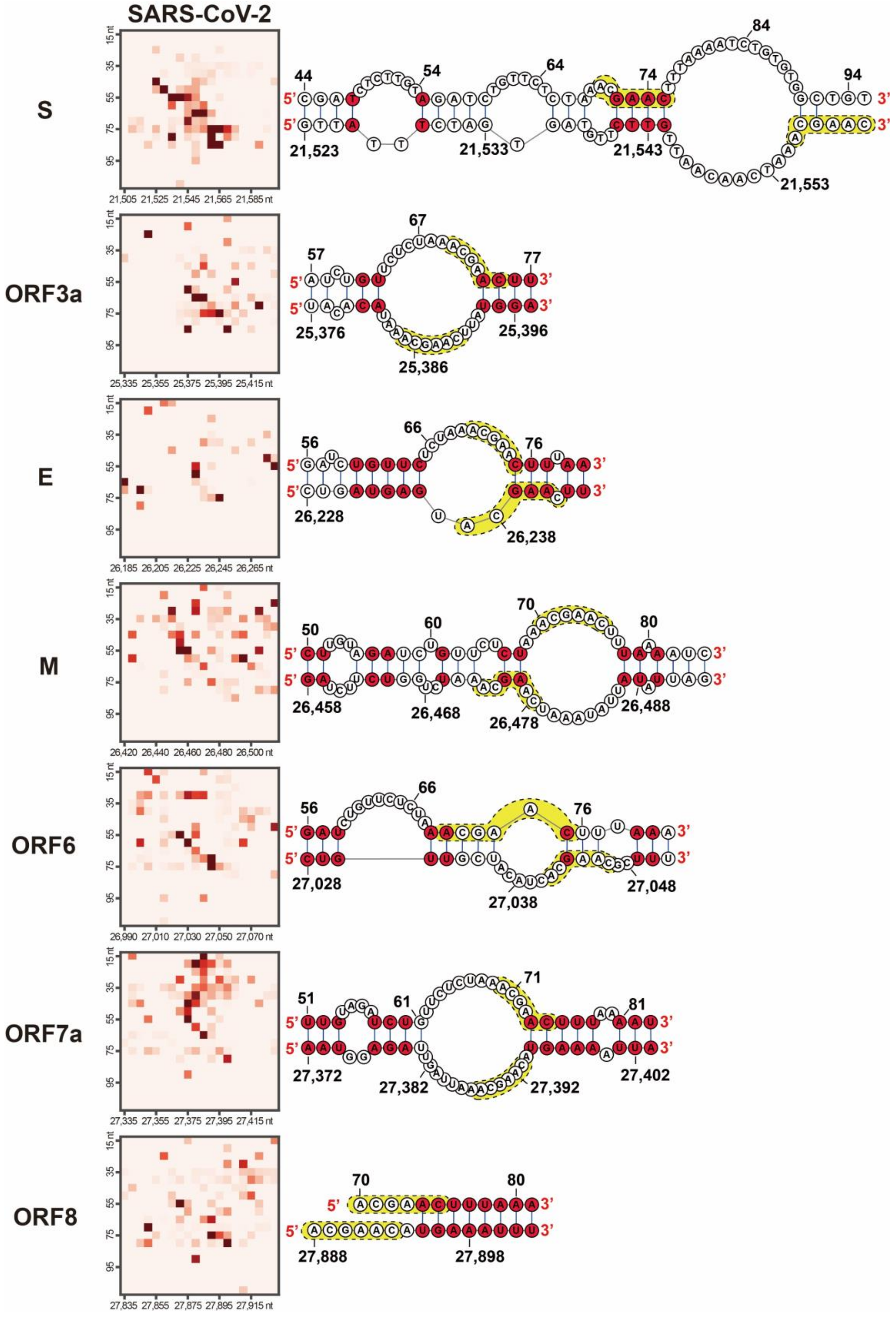


**Fig. S7 RNA-RNA interactions and base pairing between TRS-L flanking region and TRS-B flanking region for each gene in SARS-CoV-2.**

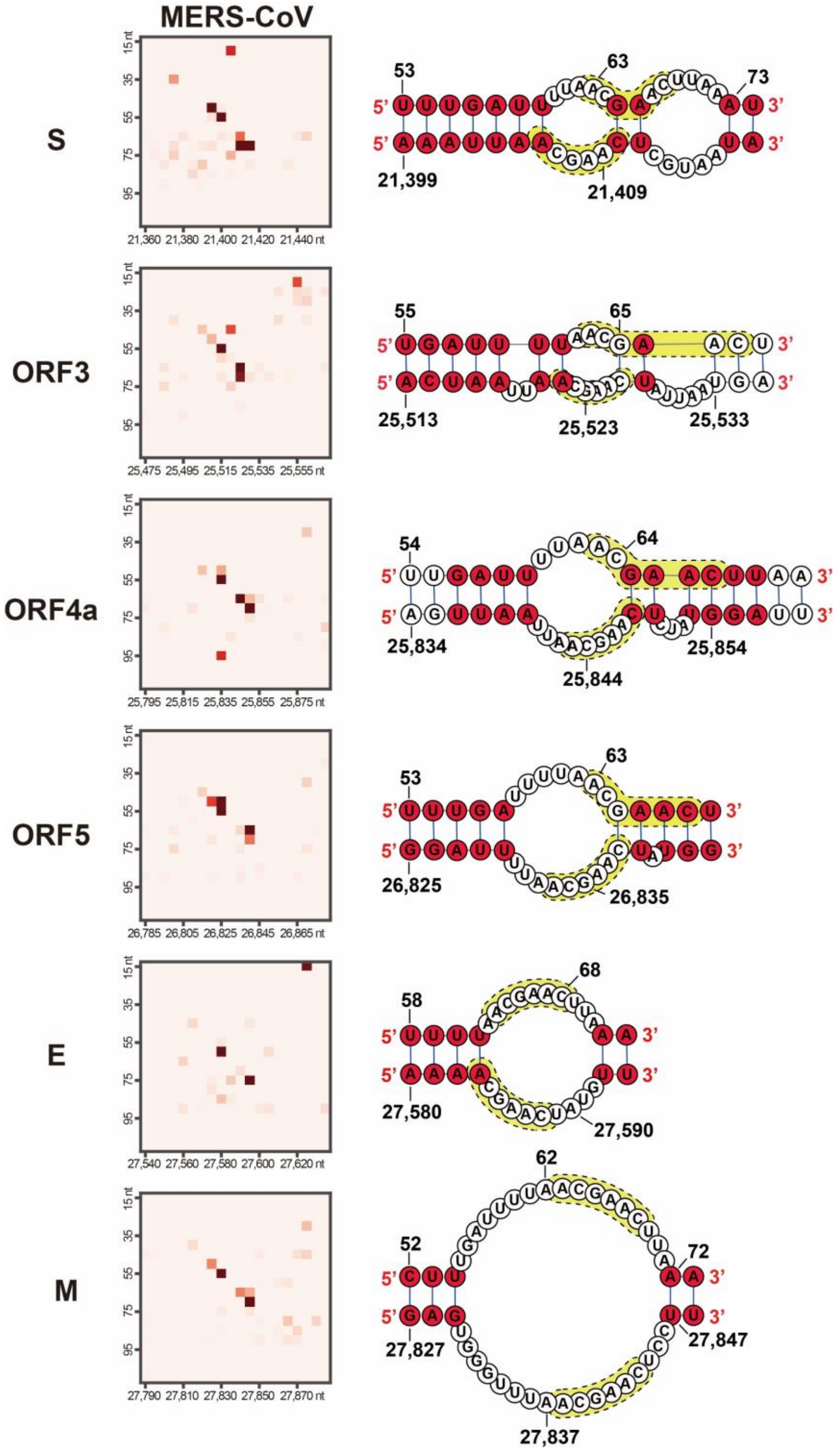


**Fig. S8 RNA-RNA interactions and base pairing between TRS-L flanking region and TRS-B flanking region for each gene in MERS-CoV.**

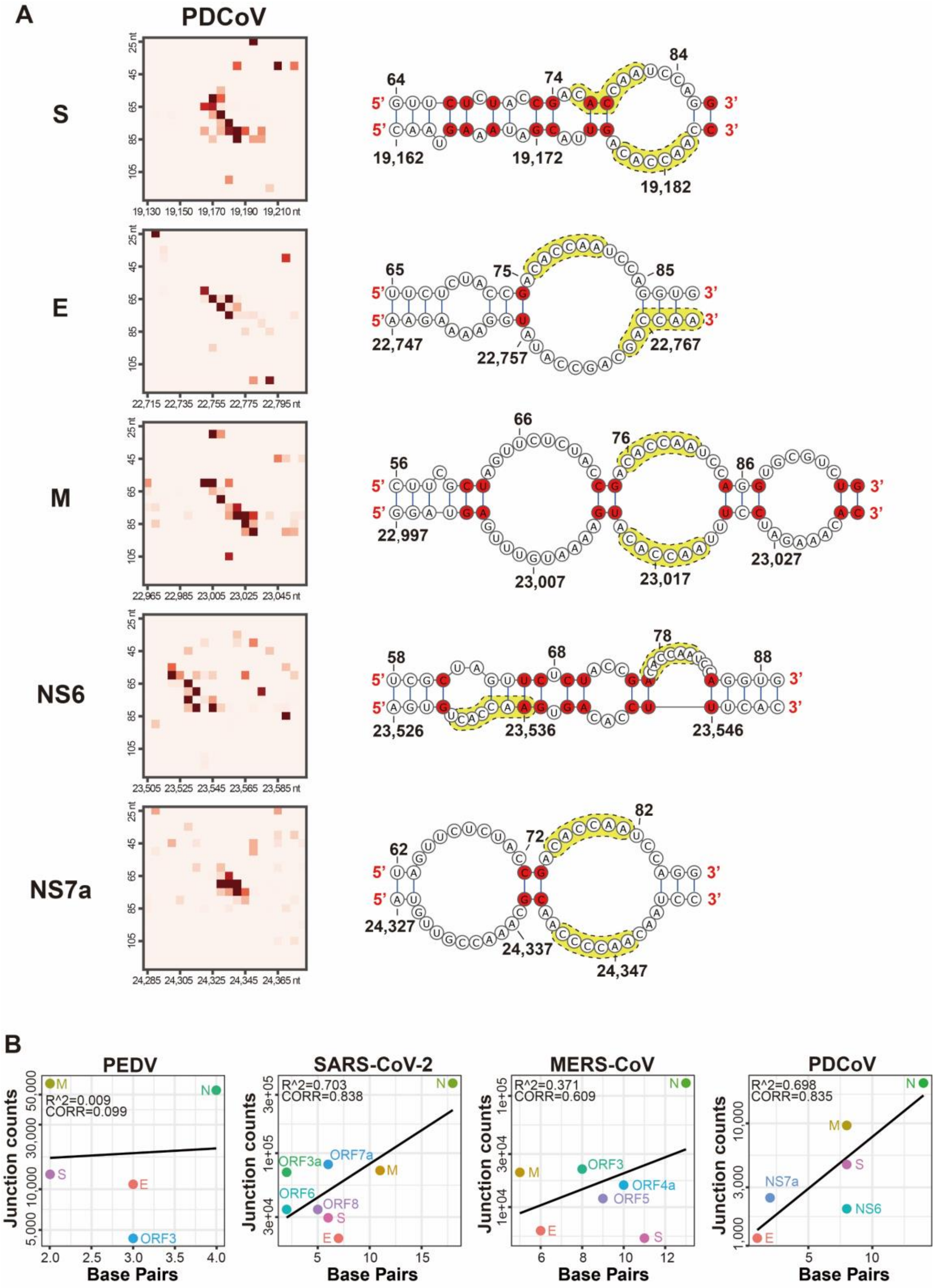


**Fig. S9 RAN secondary structures of PDCoV and correlation of RNA-seq data with RIC-seq data.**

**A**, RNA-RNA interactions and base pairing between TRS-L flanking region and TRS-B flanking region for each gene in PDCoV. **B**, Correlation between the junction counts captured by RNA-seq and the number of base pairs with interactions (highlighted in red) between TRS-B and TRS-L in the predicted secondary structures for PEDV, SARS-CoV-2, MERS-CoV and PDCoV.

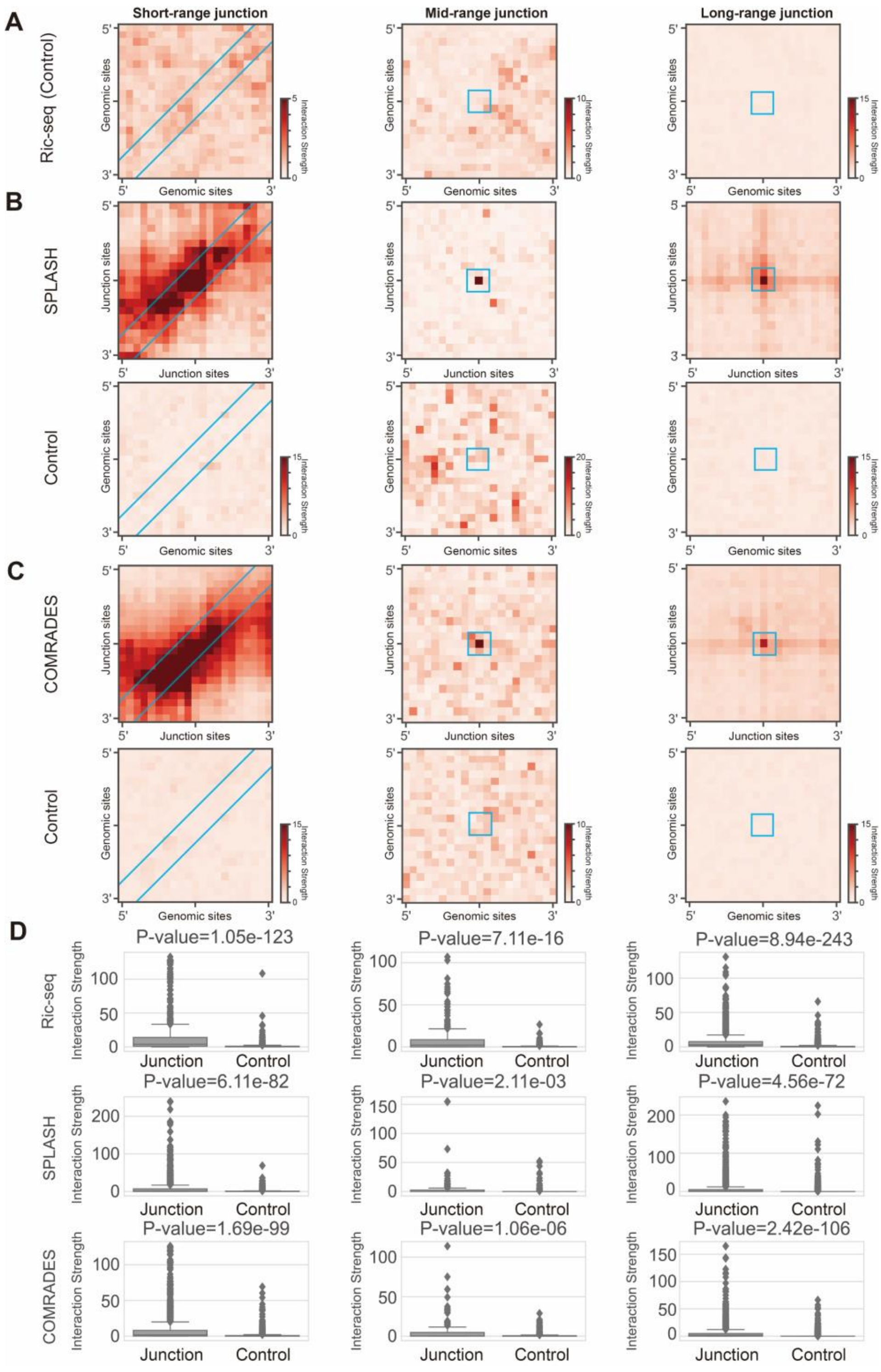


**Fig. S10 APA for non-canonical junctions of SARS-CoV-2.**

**A**, Random pairs of sequences spanning equivalent lengths as the non-canonical junctions are employed as the control. APA plot shows the aggregate RNA-RNA interactions from control groups base on RIC-seq data. **B-C**, APA plot shows the aggregate RNA-RNA interactions from non-canonical junction site-flanking regions and control groups base on SPLASH (B) and COMRADES (C) data. **D**, Boxplots of RNA-RNA interaction strengths of non-canonical junctions and control groups. Interaction strengths were determined by selecting the regions delineated between the two blue lines or blue line boxes in Fig. 4 and Fig. S10BC. The p-value of one-tailed t-test was calculated for each group.

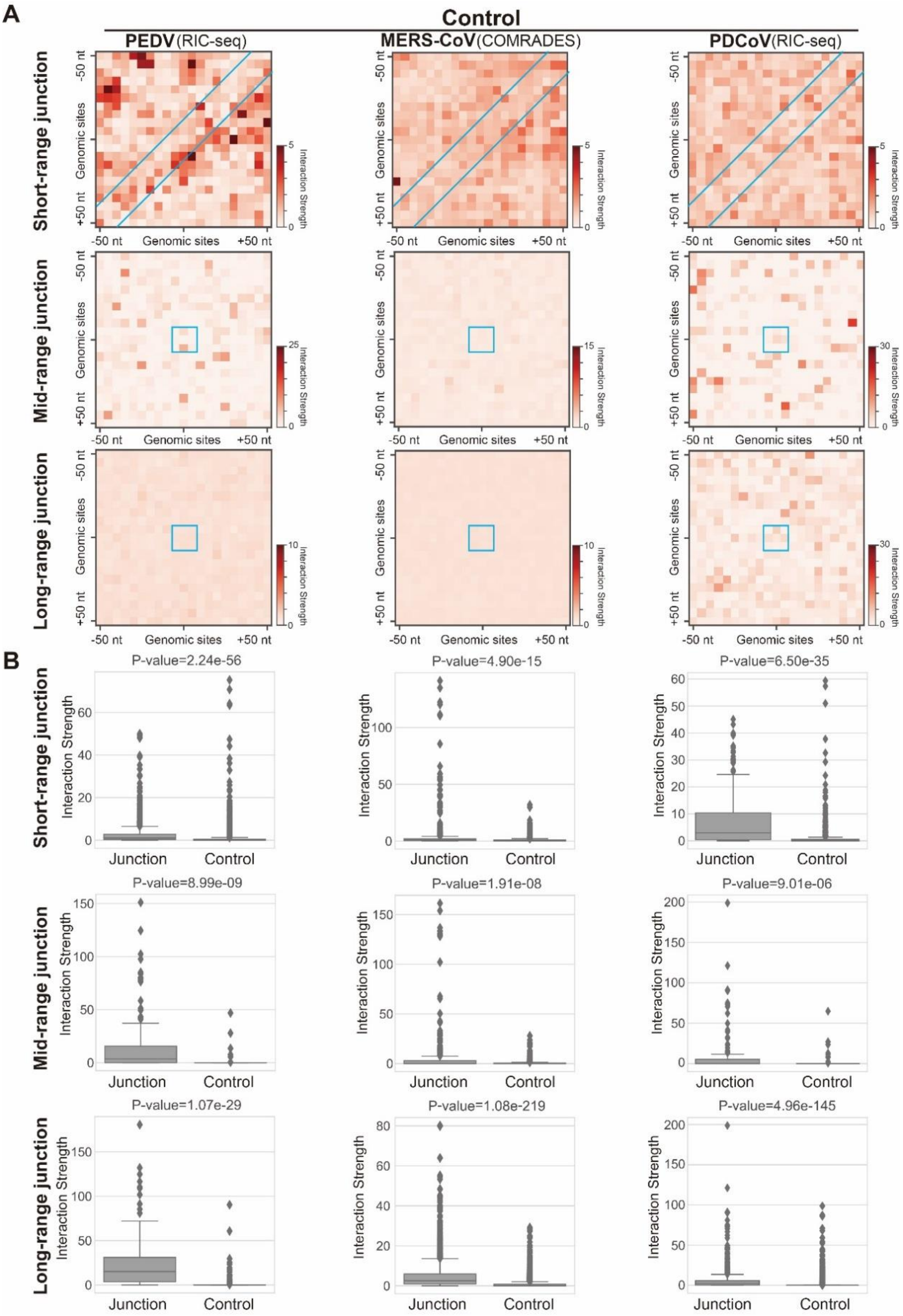


**Fig. S11 APA for non-canonical junctions of PEDV, MERS-CoV and PDCoV.**

**A**, APA plot shows the aggregate RNA-RNA interactions from control groups. **B**, Boxplots of RNA-RNA interaction strengths of non-canonical junctions and control groups. Interaction strengths were calculated by the regions located in the blue lines or blue line boxes in Fig. 4 and Fig. S11 A. The p-value of one-tailed t-test was calculated for each group.

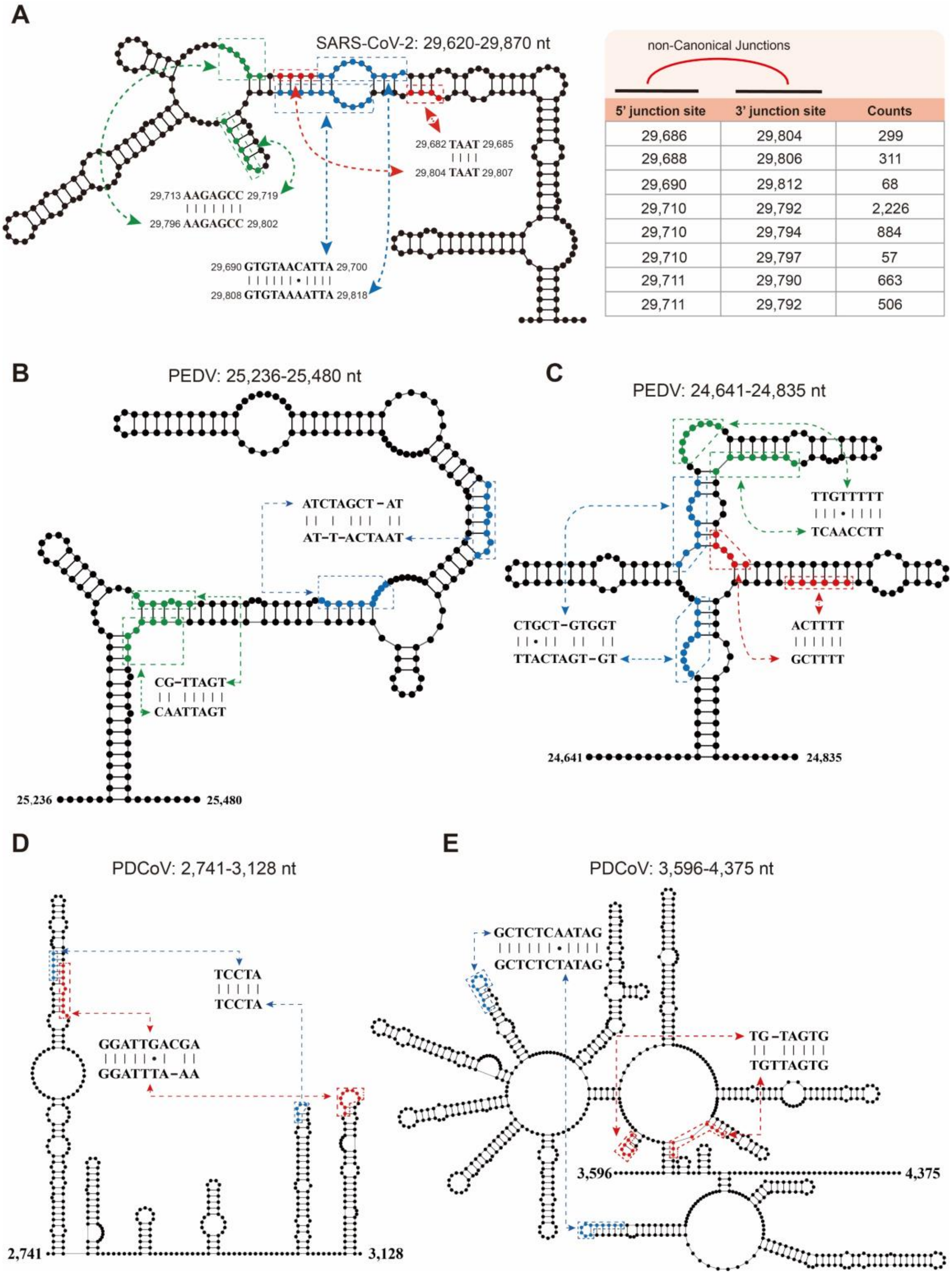


| 5' junction site | 3' junction site | Counts |
|---|---|---|
| 29,686 | 29,804 | 299 |
| 29,688 | 29,806 | 311 |
| 29,690 | 29,812 | 68 |
| 29,710 | 29,792 | 2,226 |
| 29,710 | 29,794 | 884 |
| 29,710 | 29,797 | 57 |
| 29,711 | 29,790 | 663 |
| 29,711 | 29,792 | 506 |

**Fig. S12 Stem-loops and TRS-like sequence pairs (highlighted in same color) in genomes of SARS-CoV-2 (A), PEDV (B-C) and PDCoV (D-E).**

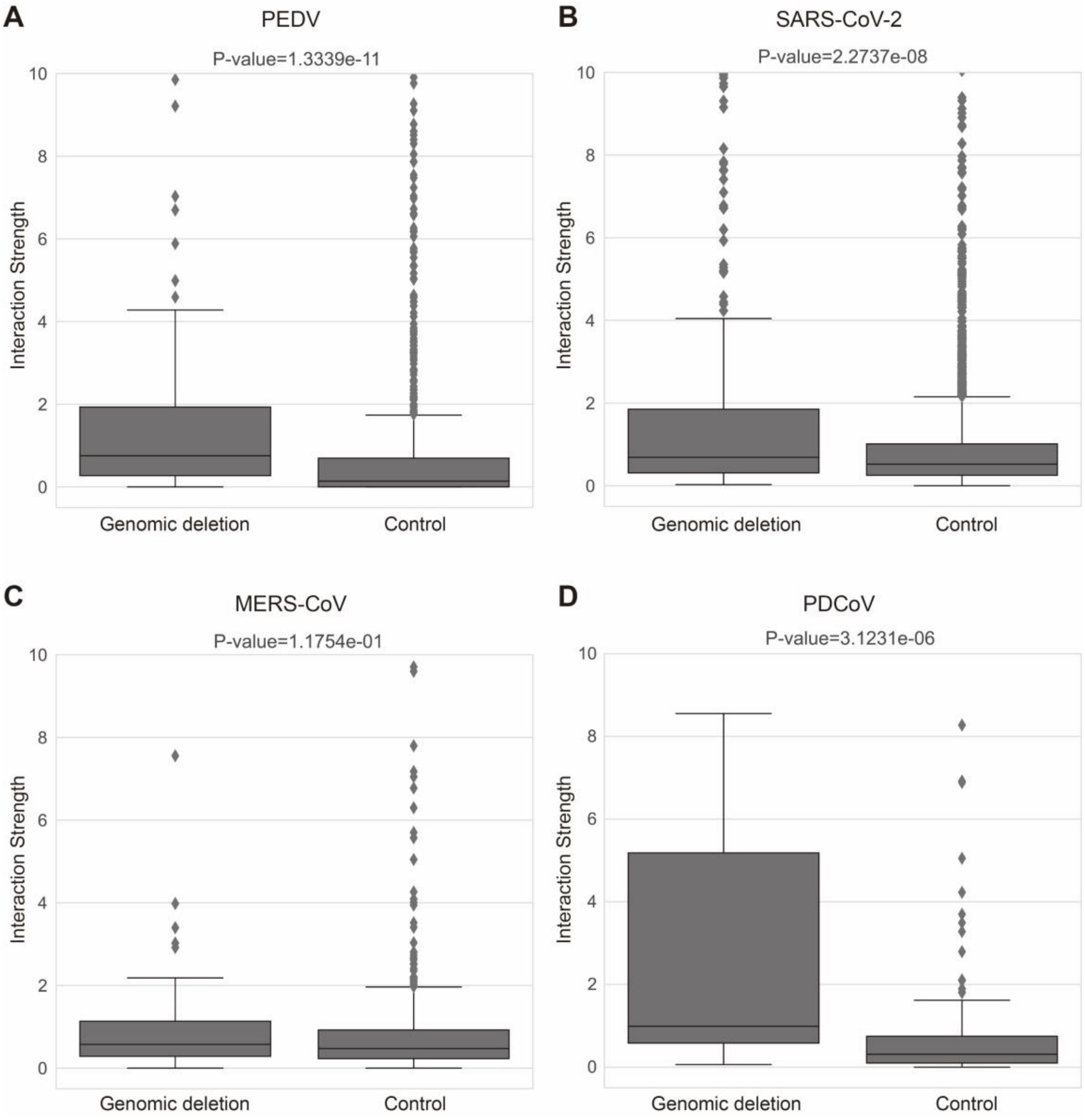


**Fig. S13 Interaction strength comparison for genomic deletions in different coronaviruses. A-D**, Boxplots depicting the RNA-RNA interaction strengths between ends of deletion regions for PEDV, SARS-CoV-2, MERS-CoV and PDCoV, compared with randomly paired genomic sites. Interaction strengths were determined by selecting the regions delineated between the two blue lines in Fig.5 A-D. The p-value of Mann-Whitney U rank test was calculated for each group.

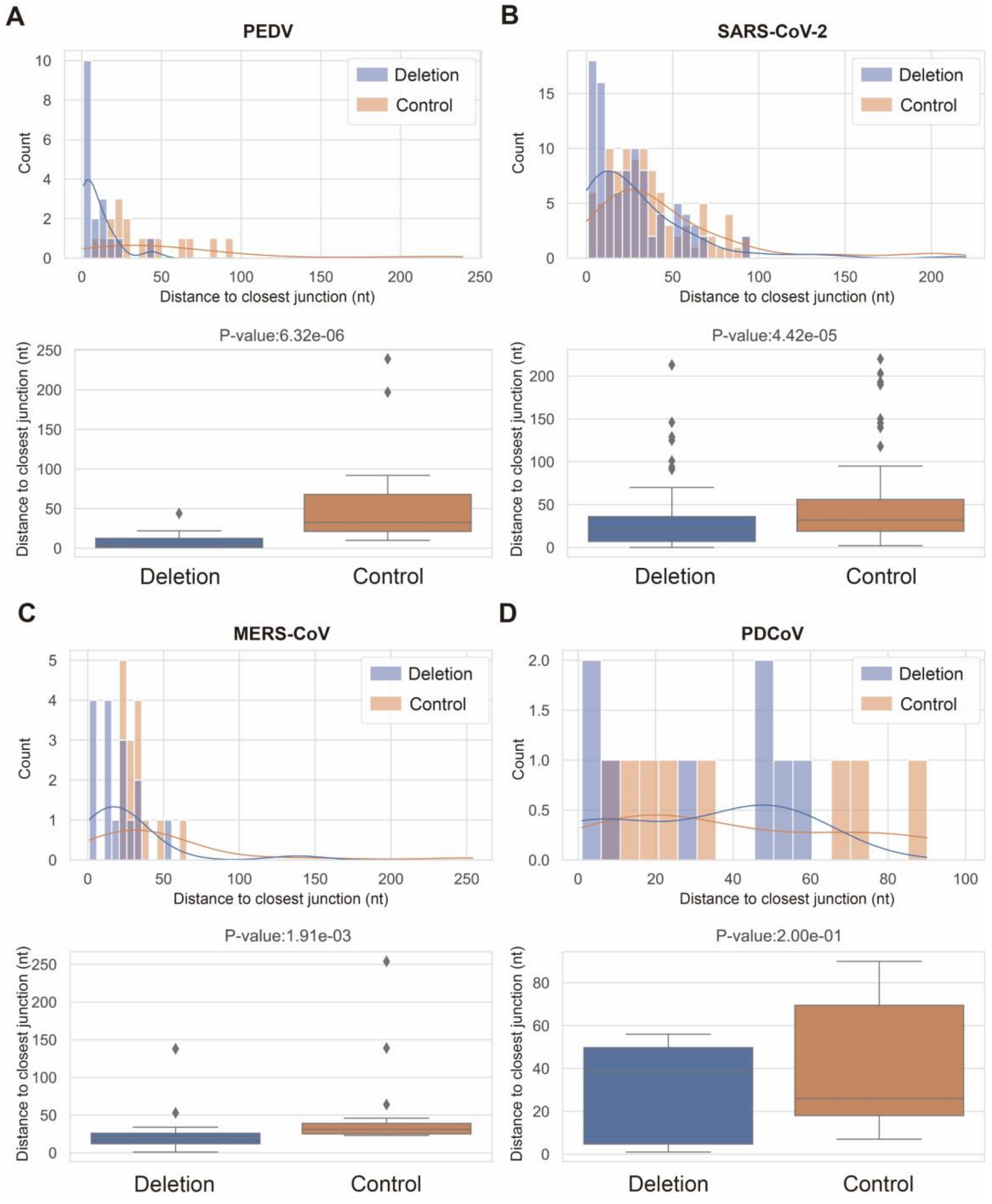


**Fig. S14 Distance between genomic deletion sites and non-canonical junction sites.**

**A**, Histogram and boxplot of the distance distribution between deletion sites and short-range junction sites for PEDV (A), SARS-CoV-2 (B), MERS-CoV (C) and PDCoV (D). The P-value of Wilcoxon rank-sum test was calculated for each group.

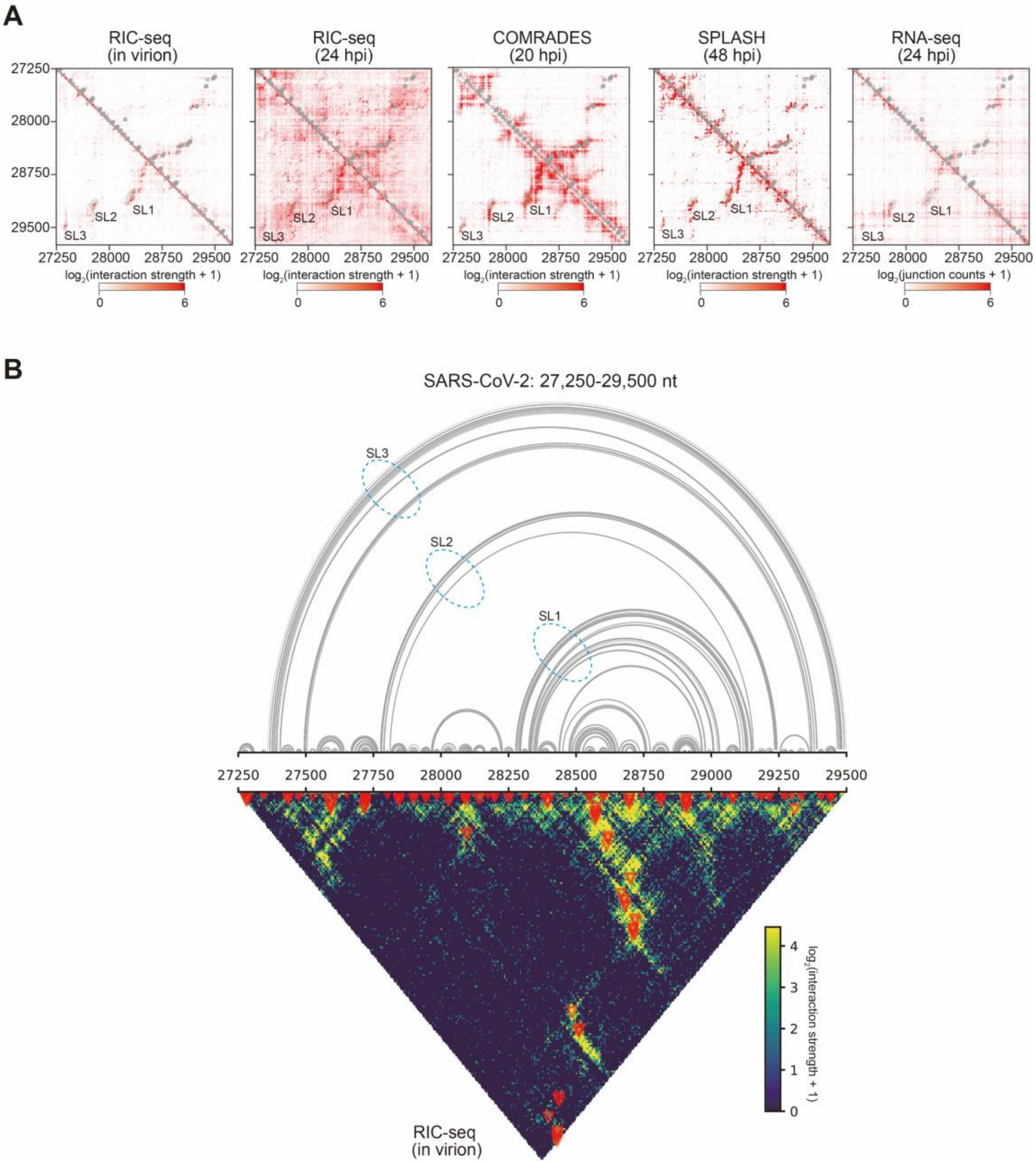


**Fig. S15 SL1/2/3 in SARS-CoV-2 genome.**

**A**, RNA-RNA interaction heatmap and short-range non-canonical junctions of SL1/2/3 regions. **B**, The base pairs within SL1/2/3 predicted from RNA-RNA interactions captured by RIC-seq (in virion).

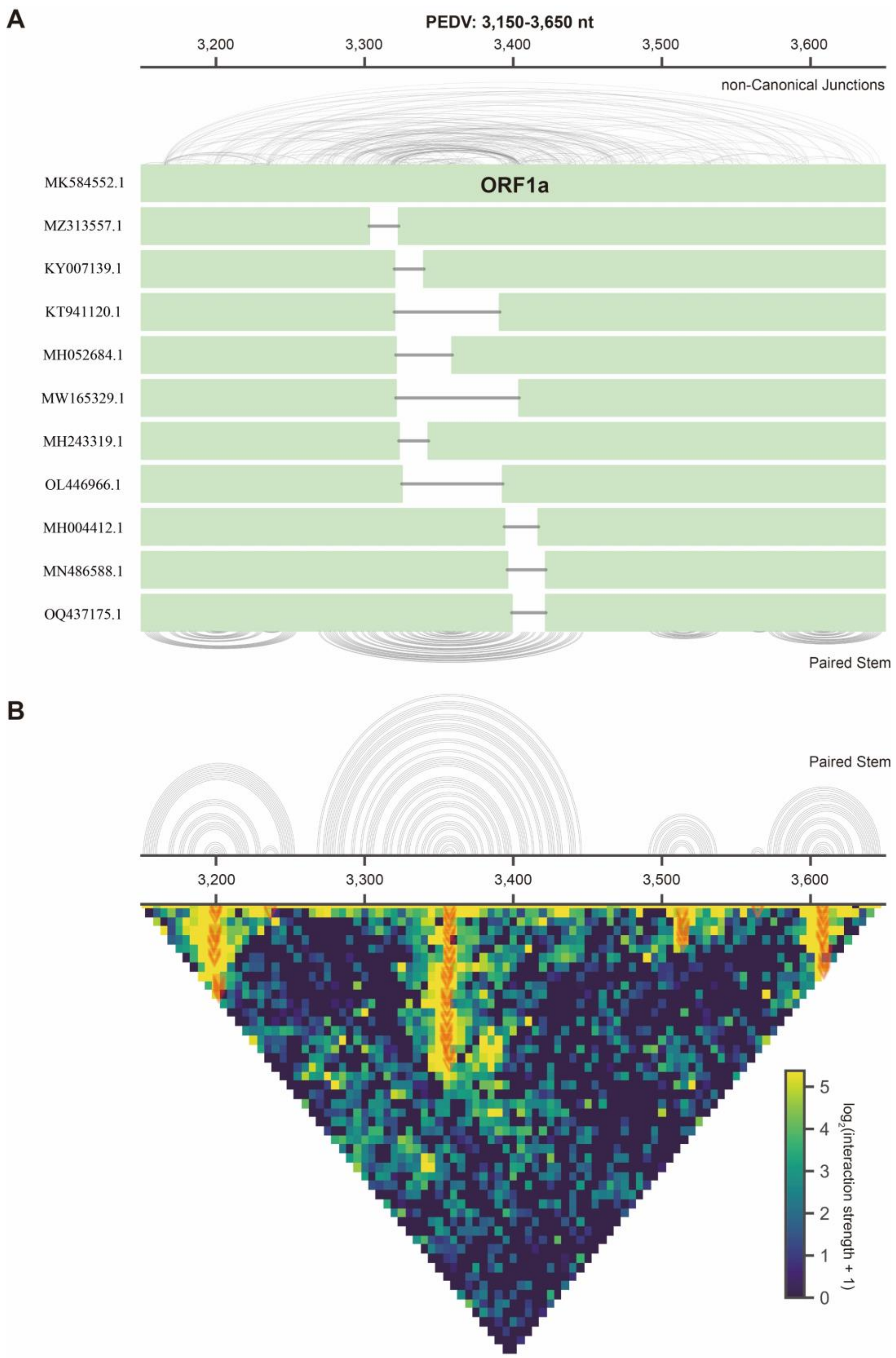


**Fig. S16 RNA-RNA interaction heatmap and secondary structure of stem-loops in PEDV genome related to genomic deletion and non-canonical junctions.**

**A**, A stem-loop region in PEDV genome (MK584552.1: 3,150-3,650 nt) contains 10 genomic deletions and abundant non-canonical junctions. **B**, RNA-RNA interaction heatmap and base pairs within the stem-loops predicted from RNA-RNA interactions captured by RIC-seq.

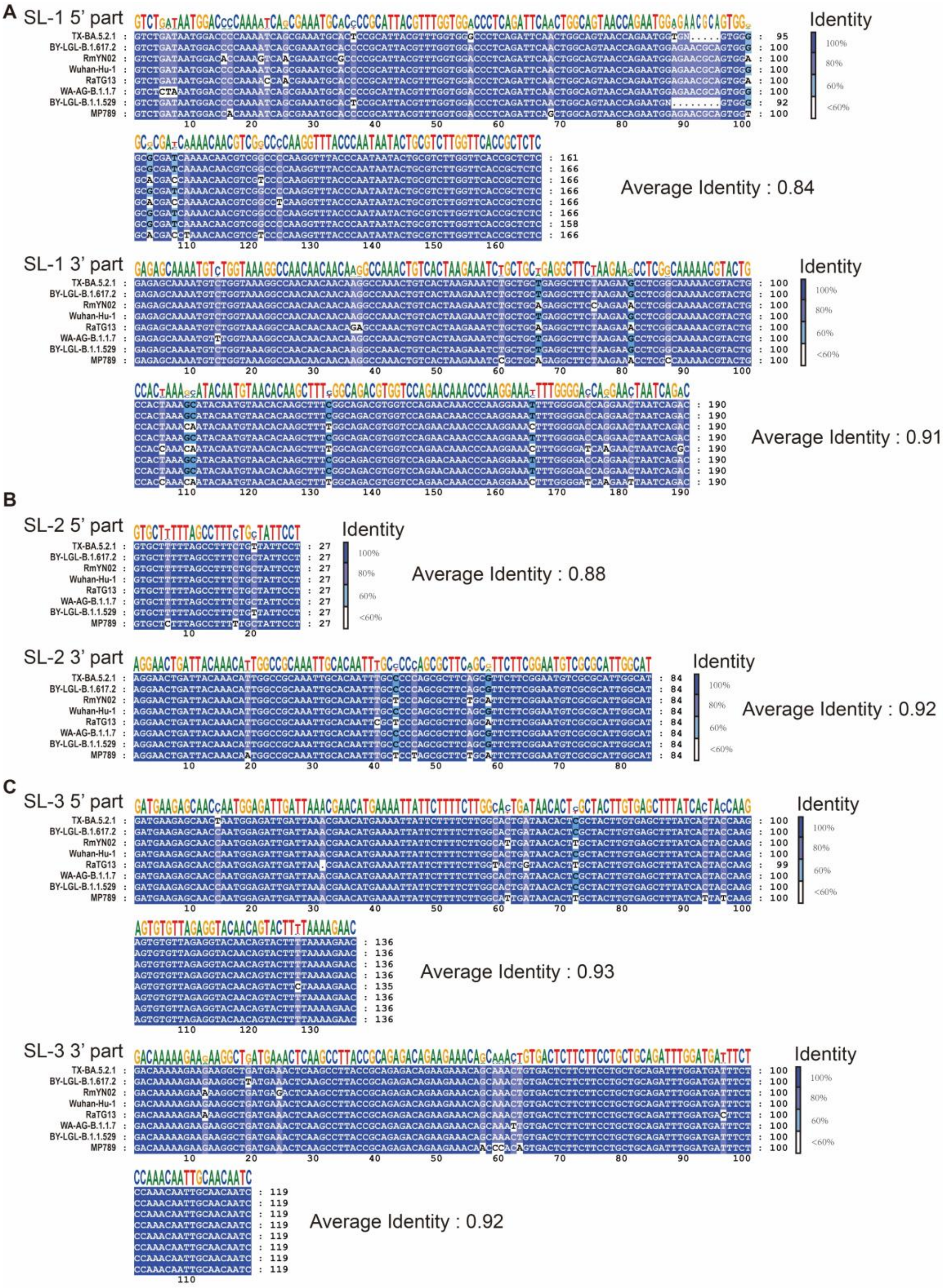


**Fig. S17 Sequence identity in the 3' and 5' parts of SL1 (A), SL2 (B), and SL3 (C) across SARS-CoV-2 related strain genomes.**

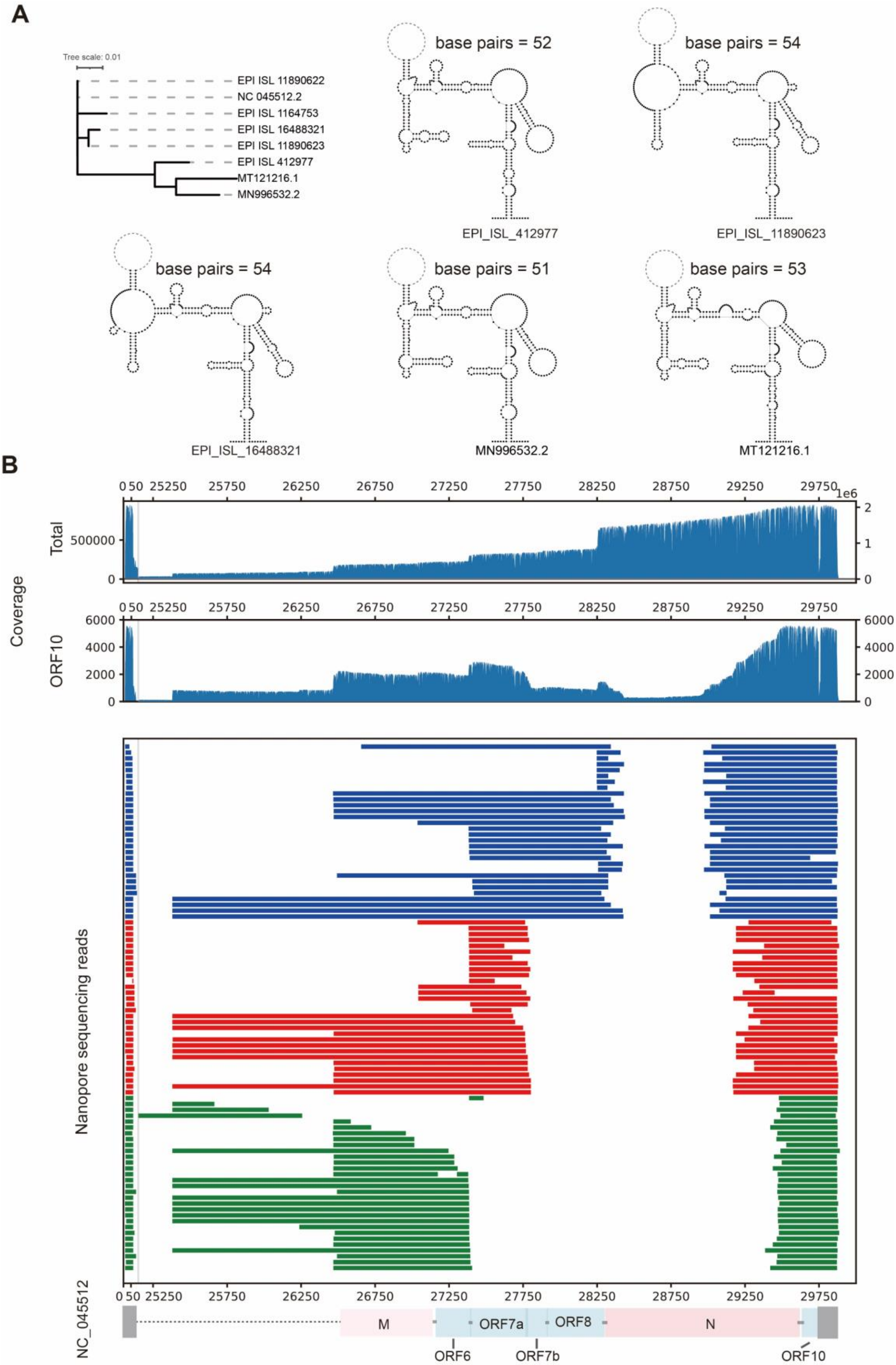


**Fig. S18 Stable SL1 and non-canonical ORF10 sgRNAs.**

**A**, Phylogenetic tree of SARS-CoV-2 related strain genomes (upper left). Secondary structure of SL1 predicted in the genome of SARS-CoV-2 related strains (lower right). **B**, Nanopore sequencing reads of ORF10 sgRNAs with double junctions.

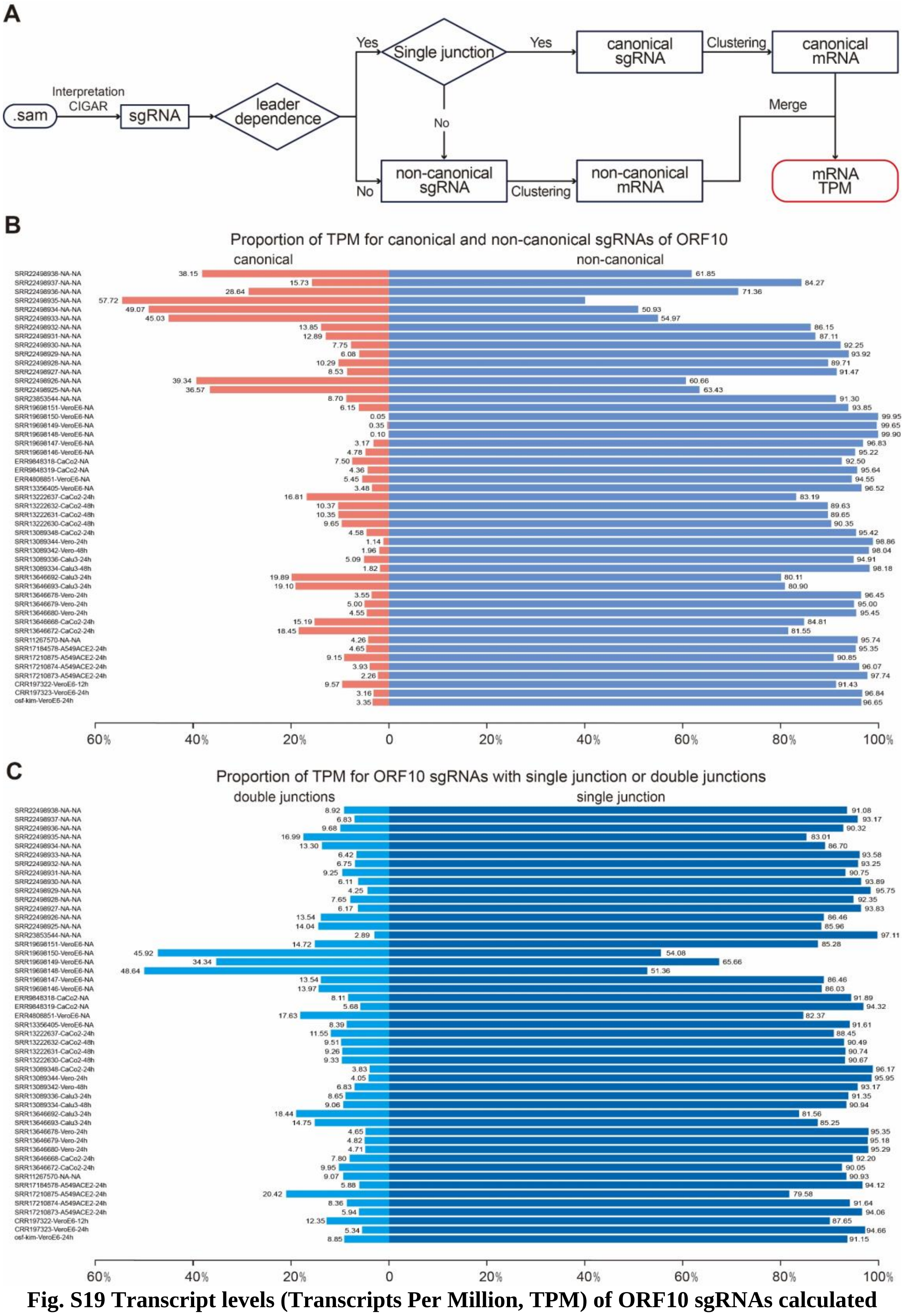


**Fig. S19 Transcript levels (Transcripts Per Million, TPM) of ORF10 sgRNAs calculated using Corona-mRNA.**

**A**, Workflow of Corona-mRNA. **B**, Proportion of TPM for canonical and non-canonical sgRNAs of ORF10 in each nanopore sample. **C**, Proportion of TPM for ORF10 sgRNAs with single or double junctions in each nanopore sample.

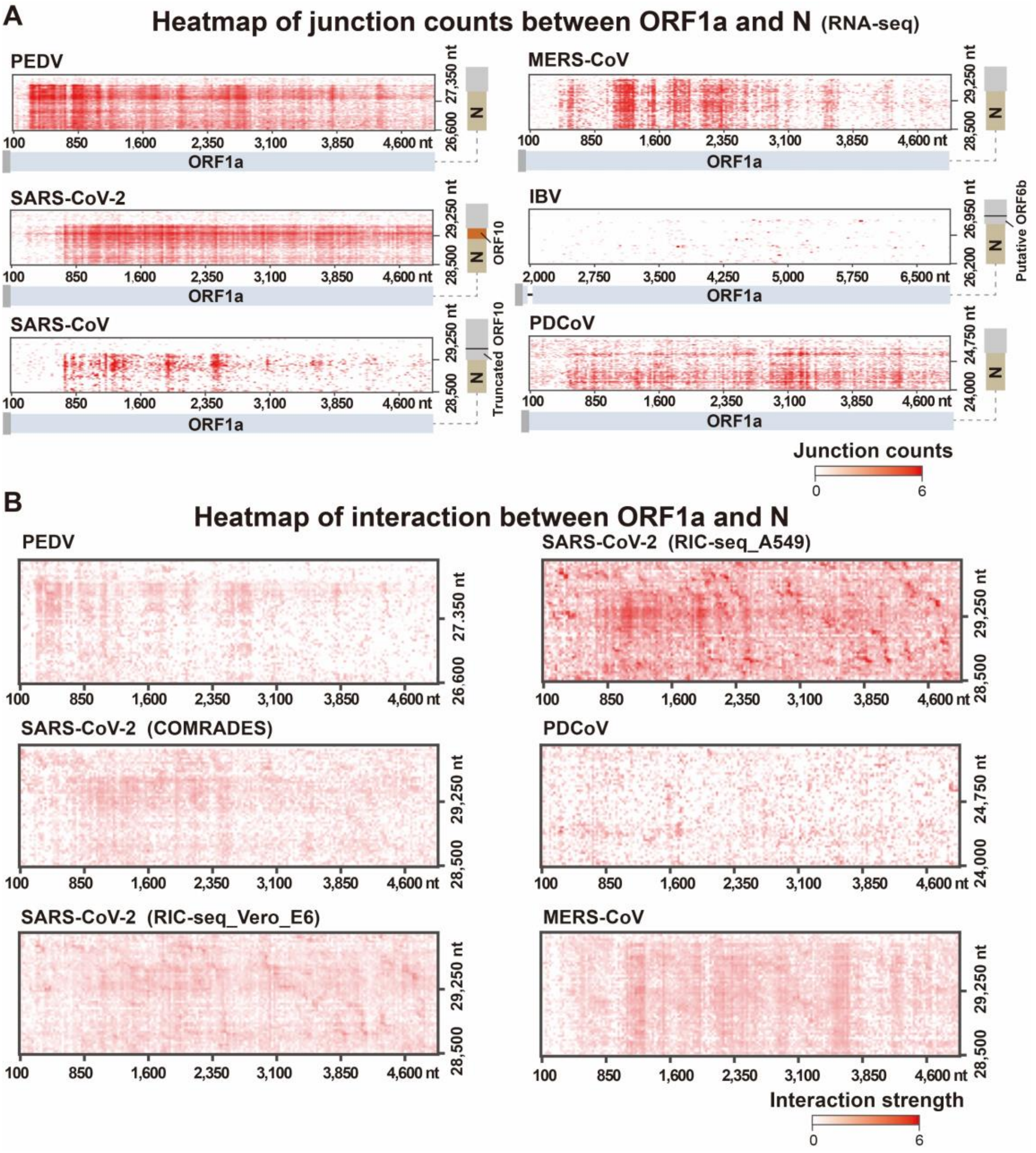


**Fig. S20 Non-canonical junctions and interactions between ORF1a and N genes in different coronavirus genomes.**

**A**, Heatmap of junction counts between ORF1a and N based on RNA-seq data from different coronaviruses. And illustrative representations of sgRNA$_{N\text{-}ORF1a}$ in the genomes of SARS-CoV-2, SARS-CoV, and other coronaviruses. **B**, Heatmap of RNA-RNA interactions between ORF1a and N based on RNA structurome data from different coronaviruses.

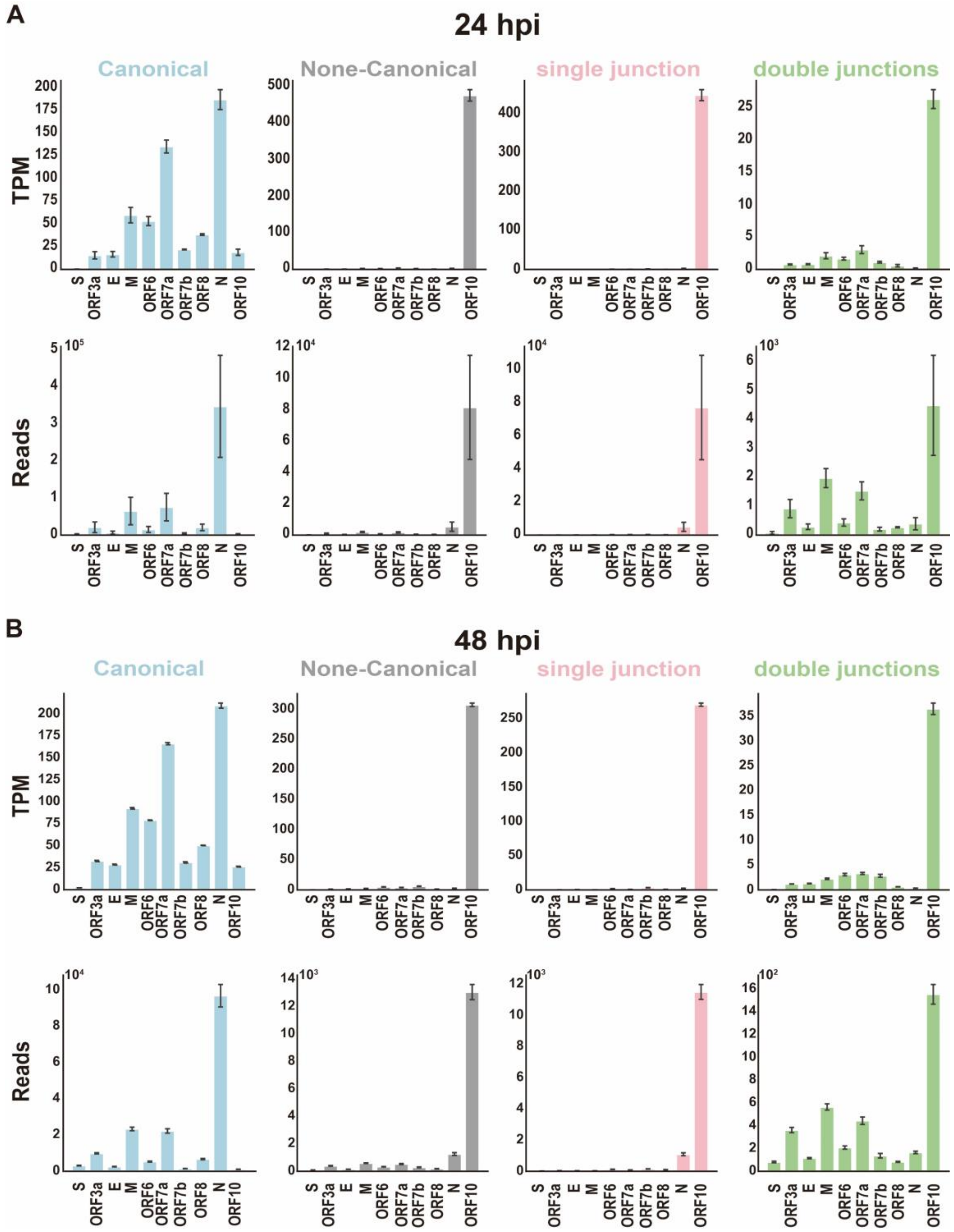


**Fig. S21 Transcription levels of individual genes in Vero E6 cells at different hpi with SARS-CoV-2.**

**A-B**, TPM and counts of reads for each gene which is transcribed through canonical and non-canonical junction at 24 hpi (A) and 48 hpi (B).

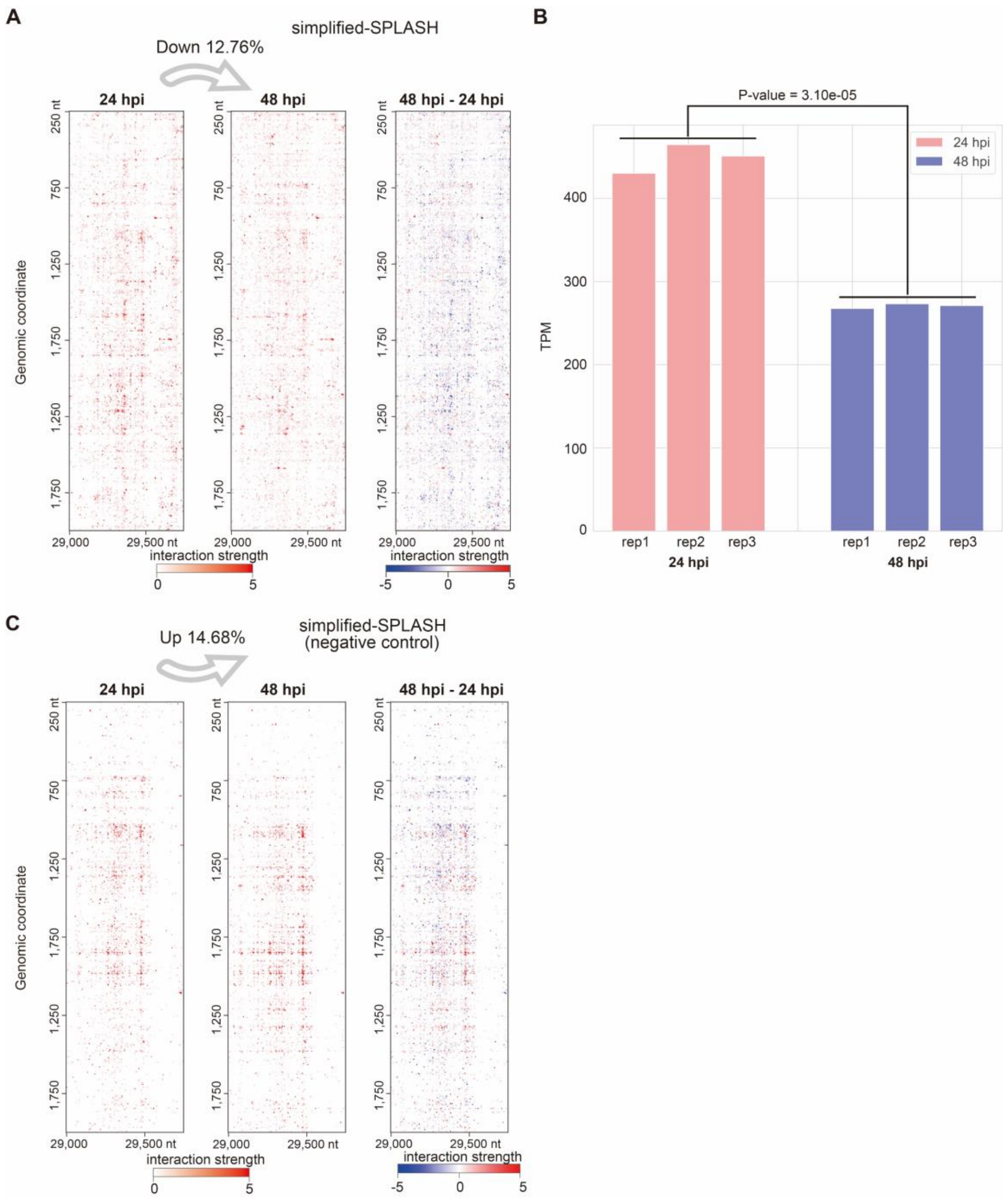


**Fig. S22 Correlation of junction counts with the RNA-RNA interaction strength.**

**A**, Alterations in RNA-RNA interactions related to ORF10 sgRNA$_{N\text{-}ORF1a}$ at 24 hpi and at 48 hpi (simplified-SPLASH). **B**, Alterations in TPM of ORF10 sgRNA$_{N\text{-}ORF1a}$ at 24 hpi and at 48 hpi. The p-value of one-tailed t-test was calculated. **C**, Alterations in RNA-RNA interactions related to ORF10 sgRNA$_{N\text{-}ORF1a}$ at 24 hpi and at 48 hpi (Negative control of simplified-SPLASH).

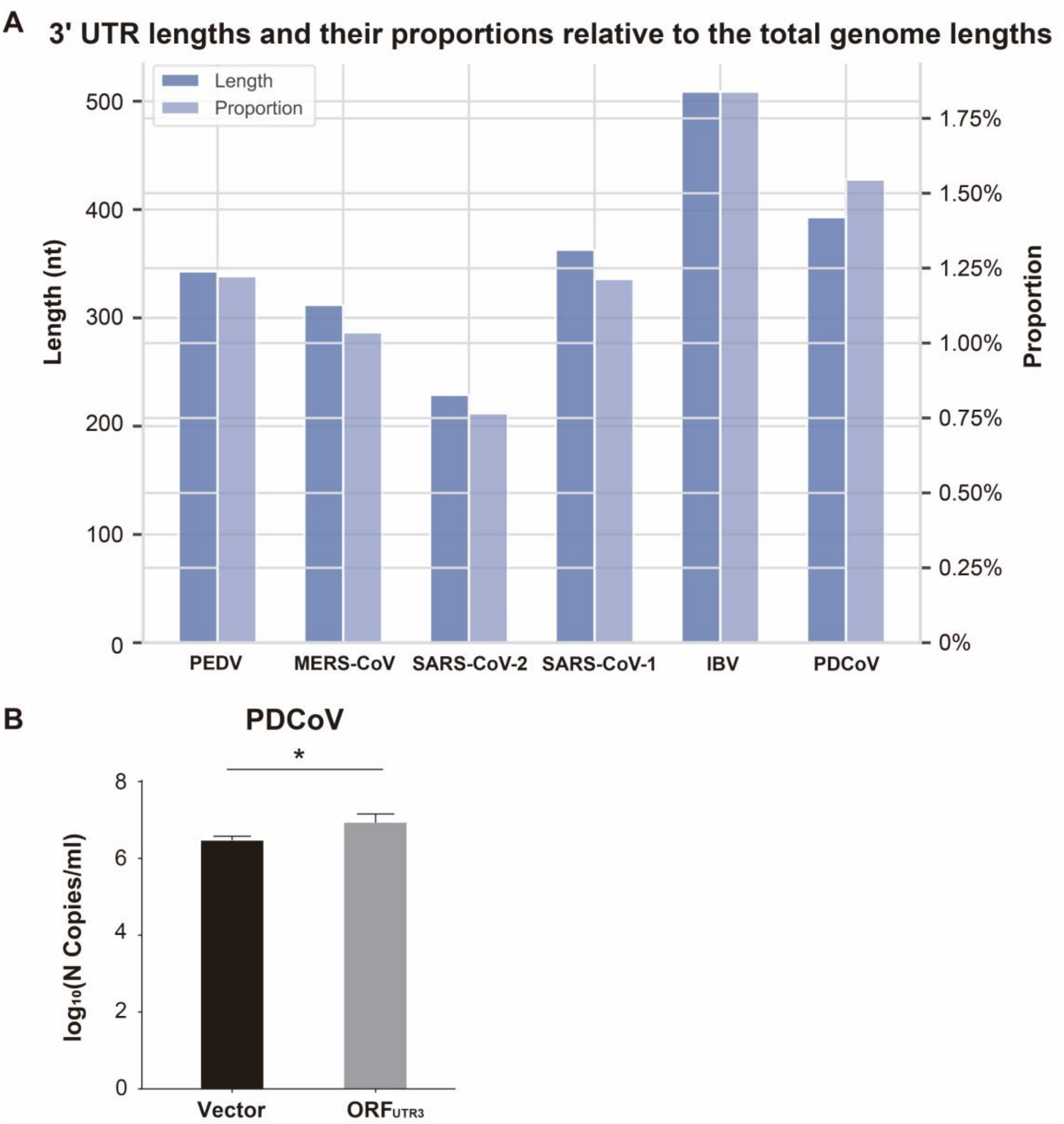


**Fig. S23 ORF$_{UTR3}$ in coronaviruses**

**A**, 3' UTR lengths and their proportions relative to the total genome lengths among different coronaviruses. **B**, LLC-PK1 cells were transfected with the ORF$_{UTR3}$ plasmid or empty vector and were then infected with PDCoV. The transcription levels N gene of PDCoV were monitored by qRT–PCR.

**TABLE S1. Proportion of non-canonical junctions at different genomic distances**

| Virus | Proportion of short-range | Proportion of mid-range | Proportion of long-range | Division distance |
|---|---|---|---|---|
| SARS-CoV-2 | 22.31% | 5.48% | 72.21% | 2,500 nt; 20,000 nt |
| MERS-CoV | 4.43% | 4.26% | 91.31% | 1,200 nt; 25,000 nt |
| PEDV | 5.39% | 2.31% | 92.30% | 1,700 nt; 22,000 nt |
| PDCoV | 48.64% | 7.49% | 43.86% | 2,100 nt; 18,750 nt |

**TABLE S2. Deletions obtained based on multiple sequence alignment**

| Virus | Number of genomes | Number of deletions | Number of deletions close to stem-loops | Proportion |
|---|---|---|---|---|
| SARS-CoV-2 | 1,110 | 105 | 31 | 29.52% |
| PEDV | 902 | 31 | 14 | 45.16% |
| MERS-CoV | 657 | 17 | 4 | 23.53% |
| PDCoV | 549 | 8 | 5 | 62.50% |

**TABLE S3. Locations of enriched genomic deletions in SARS-CoV-2 genome （NC_045512.2）**

| Reference | Location | Length | Region | Collection date | Area |
|---|---|---|---|---|---|
| EPI_ISL_414378 | 27,848-28,229 | 382 | ORF7b-ORF8 | 2020-02-17 | Singapore |
| EPI_ISL_450343 | 27,905-28,250 | 346 | ORF8 | 2020-05-09 | Bangladesh |
| EPI_ISL_832092 | 28,003-28,190 | 188 | ORF8 | 2020-04 | USA |
| EPI_ISL_426967 | 27,826-27,967 | 142 | ORF7b-ORF8 | 2020-03-27 | Australia |
| EPI_ISL_2138877 | 27,827-27,959 | 133 | ORF7b-ORF8 | 2021-05-03 | USA |
| EPI_ISL_2186398 | 27,808-27,897 | 90 | ORF7b-ORF8 | 2021-04-19 | USA |
| EPI_ISL_452497 | 27,904-27,965 | 62 | ORF8 | 2020-03-19 | Spain |
| EPI_ISL_445213 | 27,438-27,657 | 220 | ORF7a | 2020-04-28 | Bangladesh |
| EPI_ISL_1866778 | 27,504-27,689 | 186 | ORF7a | 2021-04-05 | Denmark |
| EPI_ISL_1443278 | 27,519-27,689 | 171 | ORF7a | 2021-03-18 | USA |
| EPI_ISL_2470817 | 27,519-27,725 | 207 | ORF7a | 2021-05-26 | Germany |
| EPI_ISL_806843 | 27,531-27,684 | 154 | ORF7a | 2020-12-08 | USA |
| EPI_ISL_2174719 | 27,536-27,651 | 116 | ORF7a | 2021-05-03 | Denmark |
| EPI_ISL_1235896 | 27,555-27,646 | 92 | ORF7a | 2021-03-07 | USA |
| EPI_ISL_2284503 | 27,563-27,640 | 78 | ORF7a | 2021-04-29 | Netherlands |
| EPI_ISL_2401741 | 27,621-27,670 | 50 | ORF7a | 2021-05-22 | Luxembour |

**TABLE S4 Locations of enriched genomic deletions in PEDV genome（MK584552）**

| Reference | Location | Length | Region | Collection date | Area |
|---|---|---|---|---|---|
| MZ313557.1 | 3,303-3,323 | 21 | ORF1a | 2016-04-01 | Poland |
| KY007139.1 | 3,320-3,340 | 21 | ORF1a | 2015-05-11 | China |
| KT941120.1 | 3,320-3,391 | 72 | ORF1a | 2014-10-15 | Viet Nam |
| MH052684.1 | 3,321-3,359 | 39 | ORF1a | 2017-12 | South Korea |
| MW165329.1 | 3,321-3,404 | 84 | ORF1a | 2013-07-01 | Taiwan |
| MH243319.1 | 3,323-3,343 | 21 | ORF1a | 2018-05 | South Korea |
| OL446966.1 | 3,325-3,393 | 69 | ORF1a | 2020 | China |
| MH004412.1 | 3,394-3,417 | 24 | ORF1a | 2016-07-19 | Mexico |
| MN486588.1 | 3,396-3,422 | 27 | ORF1a | 2016-02 | China |
| OQ437175.1 | 3,399-3,422 | 24 | ORF1a | 2021-09-21 | China |
